# ROP GTPase regulates polarised cell growth and cell division orientation during tissue development and organogenesis in *Marchantia polymorpha*

**DOI:** 10.1101/2023.05.23.541913

**Authors:** Hugh Mulvey, Liam Dolan

## Abstract

Cell polarity – broadly defined as the asymmetric distribution of cellular activities and or subcellular components within a cell – determines the geometry of cell growth and division during development. RHO GTPase proteins regulate the establishment of cell polarity and are conserved among eukaryotes. RHO Of Plant (ROP) proteins are a subgroup of RHO GTPases that are required for cellular morphogenesis in plants. However, how ROP proteins modulate the geometry of cell growth and division during the morphogenesis of plant tissues and organs is not well understood. To investigate how ROP proteins function during tissue development and organogenesis, we characterised the function of the single copy *ROP* gene of the liverwort *Marchantia polymorpha*. *M. polymorpha* develops morphologically complex three-dimensional tissues and organs exemplified by air chambers and gemmae, respectively. Mp*rop* loss of function mutants form defective air chambers and gemmae, indicating ROP function is required for tissue development and organogenesis. During air chamber and gemma development in wild-type, the MpROP protein is enriched to sites of polarised growth at the cell surface and accumulates at the expanding cell plate of dividing cells. Consistent with these observations, polarised cell growth is lost, and cell divisions are misoriented in Mp*rop* mutants. We propose that ROP regulates both polarised cell growth and cell division orientation in a coordinated manner to orchestrate tissue development and organogenesis in land plants.

## INTRODUCTION

Morphologically complex multicellular organisms develop three-dimensional tissues and organs. The spatial regulation of cell growth and cell division are important determinants of cellular organisation in tissues and organs. Cell polarity is one factor which influences the site and orientation of cell growth and division^1^. RHO GTPase proteins regulate cell polarity and influence the organisation of the cytoskeleton during cell growth and division in diverse eukaryotes^2^. RHO Of Plants (ROP) form a plant specific clade of RHO GTPases and ROP function is required for polarised cell growth^3^. This includes the extremely polarised tip growth of root hairs and pollen tubes^4–6^, as well as anisotropic diffuse growth of pavement cells in the leaf epidermis of *Arabidopsis thaliana*^7,8^. ROP proteins have also been implicated in orienting asymmetric cell divisions during *Zea mays* subsidiary cell differentiation in the epidermis^9^ and *Physcomitrium patens* protonema branching^10^. The ability of ROP proteins to localise to restricted domains within a cell is fundamental to their role in cell polarisation. Disruption of the spatial pattern of ROP activity, caused by the overexpression of *ROP* genes, or through loss of function mutations in genes encoding ROP regulators, result in defective polarised cell growth in *A. thaliana*^5,11–13^ and *P. patens*^14,15^. Taken together the evidence indicates that ROP activity, and its spatial regulation, are required for cell polarisation in diverse plant species.

Given the demonstrated role of ROP proteins in the generation of cell polarity, which in turn influences cell growth and division orientations, it is hypothesised these proteins function during the morphogenesis of tissues and organs in plants. Although ROP function in cellular morphogenesis is well characterised, the function of ROP proteins in tissue development and organogenesis is not well understood. This is because loss of function *rop* mutants in *A. thaliana*, the species in which ROP function has been most extensively studied, lack severe tissue and organ defects. Even in higher order *rop* mutants, which have clear defects in both pavement cell and root hair morphogenesis, the morphogenesis of shoot and root tissues and organs are similar to wild-type^16–19^. A range of tissue defects can be caused by ectopic overexpression of the dominant negative or constitutively active forms of AtROP2^20,21^. However, because the aberrant forms of AtROP2 are expressed to much higher levels and in cells where the native At*ROP2* gene is not usually expressed, it is difficult to infer ROP function in tissue development and organogenesis just from gain-of-function lines. The *P. patens rop1 2 3 4* quadruple mutant has been shown to be defective in protonema development, suggesting ROP function in tissue patterning^22^. However, the *P. patens* protonema is a relatively simple filamentous structure which grows only in two dimensions. Therefore, an understanding of how ROP function contributes to the formation of complex three-dimensional structures is still lacking.

The liverwort *Marchantia polymorpha* is an experimentally tractable system for the study of ROP function in tissue development and organogenesis. Unlike most other land plants, liverworts are unique in encoding only a single *ROP* gene in their genomes (Figure S1)^23^. The lack of severe tissue defects in *A. thaliana* loss of function *rop* mutants is likely due to functional redundancy among some of the 11 At*ROP* genes. *M. polymorpha* therefore provides a unique opportunity to take a reverse genetic approach to study *ROP* function in the absence of genetic redundancy. Furthermore, *M. polymorpha* belongs to the complex thalloid lineage of liverworts and is thus characterised by the presence of intricate three-dimensional tissue structures like the air chamber and complex organs like gemma that develop into the thallus (the haploid vegetative plant body)^24,25^. It is thus a tractable experimental system to investigate how ROP shapes complex plant tissues and organs.

To discover ROP function in plant morphogenesis, we characterised the function of the single *ROP* gene in tissue development and organogenesis in *M. polymorpha*. We show that Mp*ROP* is required for air chamber development and gemma morphogenesis. In combination with studying the cellular defects of Mp*rop* mutants, the Venus-MpROP subcellular localisation patterns demonstrate that MpROP contributes to tissue development and organogenesis by controlling anisotropic diffuse growth and cell division orientation.

## RESULTS

### Mp*ROP* is required for tissue development and organogenesis

To investigate the function of ROP signalling in the morphogenesis of complex plant structures, we set out to define the function of the single Mp*ROP* gene during the vegetative development of *M. polymorpha*. The dorsal (upper) surface of the *M. polymorpha* thallus (Figure 1A, E) comprises the air chamber tissue (Figure 1I) and is also where gemma cups develop (Figure 1M) in which asexual propagules called gemmae are produced (Figure 1W). The Mp*ROP* gene is expressed ubiquitously during the formation of air chambers and gemmae (Figure S2). To determine if Mp*ROP* is required for the morphogenesis of these structures, we generated 21 independent Mp*rop* mutant alleles using CRISPR/Cas9 mediated mutagenesis and selected two complete loss of function alleles (Mp*rop-1* and Mp*rop-3*) and one partial loss of function allele (Mp*rop-2*) for phenotypic analysis (Table S1, see Figure S3 for details of the three alleles).

**Figure 1.**
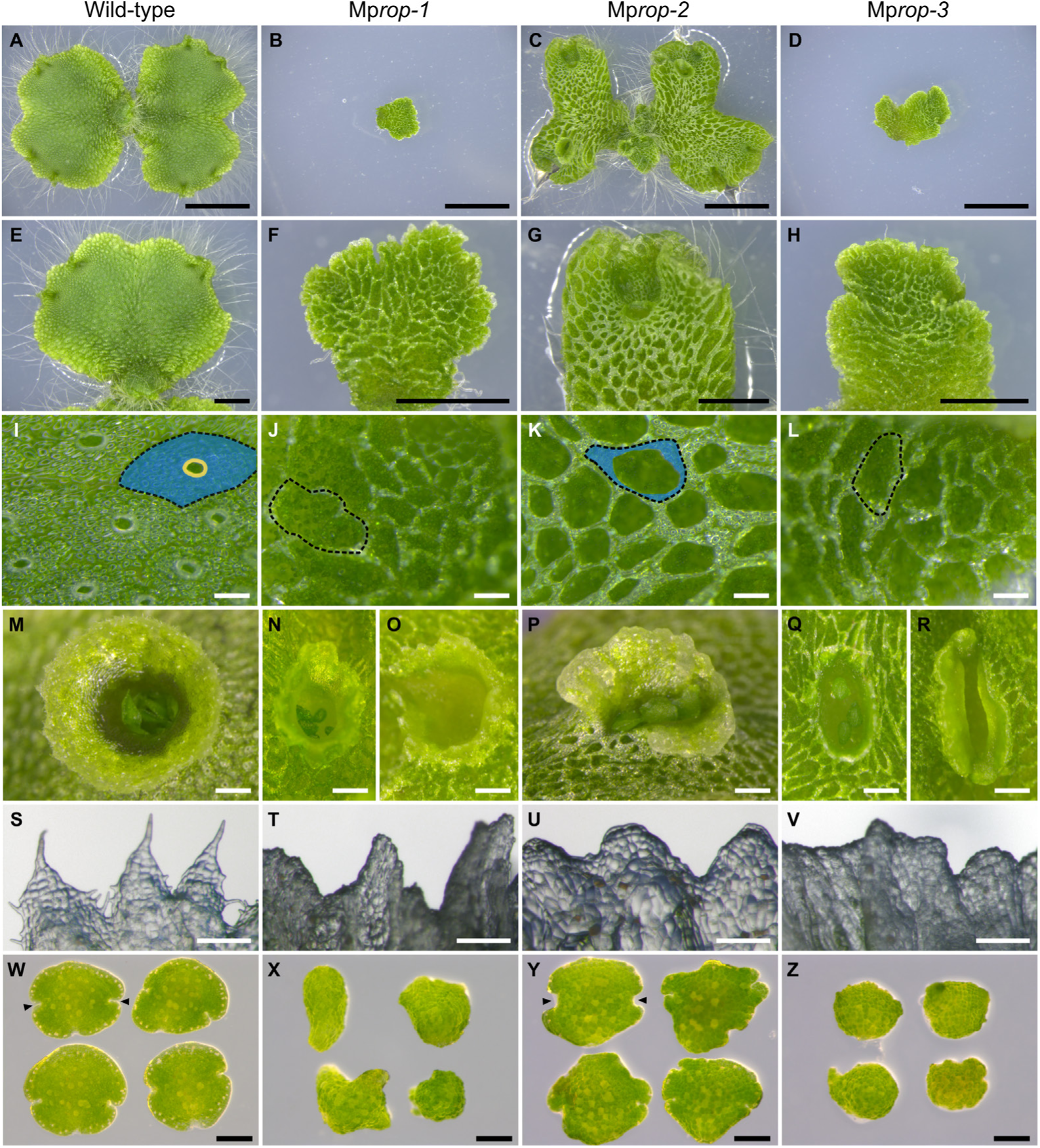
Mp*ROP* is required for tissue development and organogenesis. Stereomicroscope images of (A-L) the dorsal thallus surface of 14-day old gemmalings, (M-V) gemma cups, and (W-Z) 0-day old gemmae of wild-type, Mp*rop-1*, Mp*rop-2*, and Mp*rop-3*. (A-D) Whole gemmaling (scale bar, 5mm). (E-H) Single thallus lobe (scale bar, 2mm). (I-L) Dorsal thallus epidermis (scale bar, 200μm). The boundary of one air chamber is demarcated with a black dashed line. The air chamber roof is highlighted in blue and the outline of an air pore is marked in yellow. In Mp*rop-1* and Mp*rop-3*, air chamber boundaries are visible, but air chamber roofs and the air pores are missing. In Mp*rop-2*, air chamber roofs are partially present, but air pores are absent. Consequently, in Mp*rop-1*, Mp*rop-2*, and Mp*rop-3*, the subepidermal floor of the air chambers is visible through the enlarged aperture, unlike in wild-type. (M-R) Gemma cup (scale bar, 500μm). Mp*rop* mutant gemma cups are irregular in structure and Mp*rop-1* (N, O) and Mp*rop-3* (Q, R) gemma cups contain very few or no gemmae. (S-V) Gemma cup rim (scale bar, 200μm). Wild-type cup rim is characterised by an intricate serration pattern (S). This is lost in Mp*rop* mutants. (W-Z) Gemmae (scale bar, 200μm). Mp*rop* mutant gemmae are morphologically defective. Arrowheads point to meristematic notches. Mp*rop-1* and Mp*rop-3* gemmae lack recognisable meristematic notches.

In contrast to wild-type air chambers – which comprise an epidermal roof with a central air pore (in blue and yellow, respectively in Figure 1I) covering an intercellular air space and a subepidermal chamber floor – air chambers of the complete loss of function mutants, Mp*rop-1* and Mp*rop-3*, lack the epidermal roof and the air pore (Figure 1J, L). Consequently, walls separating neighbouring air chambers and the subepidermal chamber floors are exposed. In the partial loss of function mutant, Mp*rop-2*, air pores are also absent and a partial roof forms (Figure 1K). Therefore, Mp*ROP* is required for the formation of the air chamber roof and the air pore.

Mp*ROP* is also required for gemma cup and gemma morphogenesis. Wild-type gemma cups are radially symmetric structures (Figure 1M) with an intricately serrated rim (Figure 1S). By contrast, Mp*rop* mutant gemma cups are irregular in structure (Figure 1N-R) and lack an intricately serrated rim (Figure 1T-V), demonstrating that Mp*ROP* is required for gemma cup morphogenesis. Wild-type gemmae are disk-shaped – the gemma edge is uniformly curved with two lateral apical notches, each housing a meristem (Figure 1W). By contrast, gemmae of the complete loss of function mutants, Mp*rop-1* and Mp*rop-3*, are globular and lack recognisable notches (Figure 1X, Z), whilst some gemmae of the partial loss of function mutant, Mp*rop-2*, have an uneven edge (Figure 1Y). Moreover, Mp*rop-1* and Mp*rop-3* produce fewer gemmae than wild-type, with some mutant gemma cups being empty (Figure 1O, R). These phenotypes indicate that Mp*ROP* function is required for both the initiation of gemma formation at the base of the gemma cup and for gemma morphogenesis. Introduction of a wild-type copy of Mp*ROP* in both Mp*rop-1* and Mp*rop-2* restored wild-type like morphology, confirming that the observed mutant phenotypes are due to a loss of Mp*ROP* function (Figure S4). Taken together, the defective air chambers, gemma cups and gemmae in Mp*rop* mutants demonstrate that Mp*ROP* is required for the morphogenesis of complex structures derived from the dorsal epidermis.

To determine how MpROP function contributes to organogenesis and tissue development, we investigated MpROP function in gemma organogenesis and then in air chamber tissue development. During the morphogenesis of both structures, we examined the subcellular localisation pattern of Venus-MpROP in wild-type and then the cellular level defects in Mp*rop* mutants, to develop a spatiotemporal understanding of cellular processes regulated by MpROP.

### Venus-MpROP subcellular localisation patterns suggest MpROP regulates polarised cell growth and cell division during gemma morphogenesis

Wild-type gemma primordia, which originate from a single epidermal cell in the base of a gemma cup, initially develop as a two-dimensional, flattened, organ because the orientation of cell expansion and cell division are restricted primarily to a single plane (Figure 2A)^26,27^. The two flattened surfaces of a gemma primordium correspond to the future dorsal and ventral surfaces of the vegetative plant body that develops from the gemma. Since localised accumulation of MpROP promotes tip growth of rhizoids (Figure S5), we hypothesized that localised accumulation of MpROP promotes the anisotropic diffuse growth of epidermal cells in an orientation perpendicular to the dorsoventral axis of the gemma primordia, for its two-dimensional development. To test this hypothesis, we first determined the localisation of Venus-MpROP fusion protein in epidermal cells at the gemma primordium edge, where the future dorsal and ventral surfaces meet. Venus-MpROP is polar localised to the gemma edge in some epidermal cells (Figure 2B-D). Compared to the integral plasma membrane marker mScarletI-AtLTI6b, Venus-MpROP was enriched in an externally facing cortical domain, which is roughly parallel to the dorsoventral axis (Figure 2D, E). As Venus-MpROP marks the site of polarised growth in rhizoids (Figure S5), this polar localisation at the edge of the gemma primordium is consistent with the hypothesis that MpROP promotes anisotropic diffuse growth of epidermal cells in an orientation perpendicular to the dorsoventral axis.

**Figure 2.**
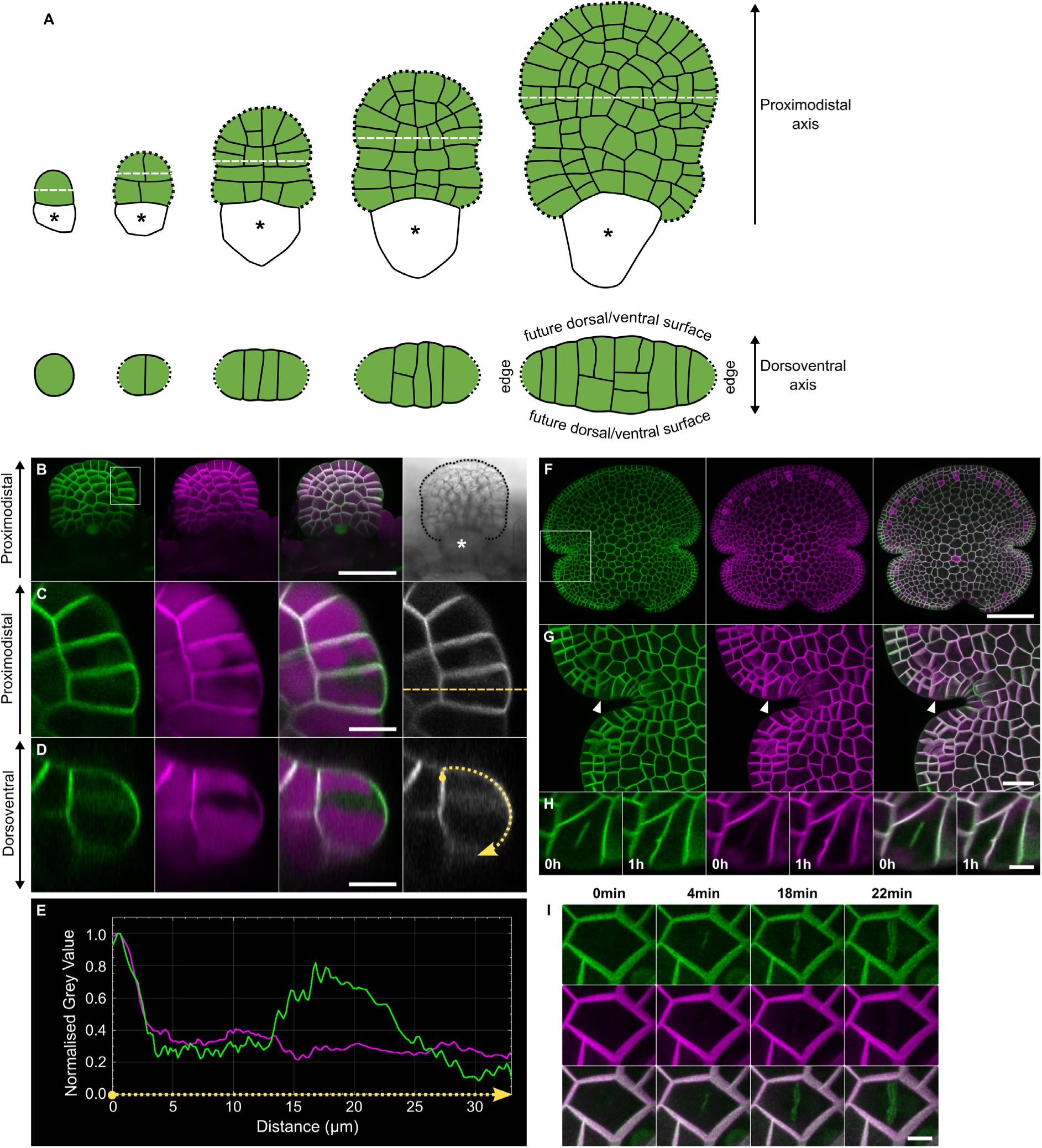
Venus-MpROP polar localised in epidermal cells during gemma morphogenesis. (A) Schematic representation of wild-type gemma primordium development. Top row shows surface view and bottom row shows transverse cross section (along white dashed line). Black dotted line marks the gemma primordium edge and asterisk marks stalk cell which connects the primordium to the gemma cup floor. Schematic based on confocal images of wild-type gemma primordia. (B) Gemma primordium developing at the base of a gemma cup, expressing *_pro_*Mp*ROP:Venus-*Mp*ROP*, *_pro_*Mp*UBE2:mScarletI*-At*LTI6b*, and *_pro_*Mp*RSL3:Venus-NLS*. From left to right, images of Venus fluoresce, mScarletI fluorescence, merged, and bright-field. Asterisk marks stalk cell. (C) Epidermal cells at the gemma primordium edge (region marked with white box in B) (D) Optical cross section (in the plane marked by the dashed yellow line in C) reveals Venus-MpROP polarised to the outer face of the epidermal cell, roughly parallel to the dorsoventral axis. (E) Normalised fluorescence intensities of Venus-MpROP (green) and mScarlet-AtLTI6b (magenta) along the plasma membrane region marked with the dotted yellow line in D. (F) Immature gemma expressing *_pro_*Mp*ROP:Venus-*Mp*ROP*, *_pro_*Mp*UBE2:mScarletI*-At*LTI6b*, and *_pro_*Mp*RSL3:Venus-NLS*. From left to right, images of Venus fluoresce, mScarletI fluorescence and merged. Absence of *_pro_*Mp*RSL3:Venus-NLS* signal indicates rhizoids initials have not yet differentiated. (G) Notch region of immature gemma (region marked with white box in F). Arrowhead marks a dividing cell. (H) Epidermal cell (marked by arrowhead in G) during division and 1 hour after division. (I) Centrifugal expansion of the Venus-MpROP (green) at the division plane of a dividing gemma epidermal cell. Scale bars, 40μm (B), 10μm (C and D), 100μm (F), 20μm (G), 5μm (H and I). Fluorescent images in B, F, G, and I are maximum Z-projection images whilst those in C, D, and H are sum projection of five consecutive images.

As well as promoting anisotropic diffuse growth, we hypothesised that MpROP regulates cell division orientation during gemma morphogenesis, because ROP-related proteins in other organisms regulate cell division orientation^10,28^. To test this hypothesis, we determined Venus-MpROP subcellular localisation in dividing epidermal cells by performing time-lapse imaging of developing gemmae. In an immature gemma, where meristematic notches have formed but rhizoid initials have not yet differentiated, Venus-MpROP was detected at the division plane of dividing cells near the meristem (Figure 2F-H). Venus-MpROP initially occupies only the central part of the division plane and later occupies the entire division plane. Finer resolution time lapse imaging revealed centrifugal expansion of the Venus-MpROP domain, consistent with the hypothesis that MpROP is localised to the expanding cell plate (Figure 2I). The plasma membrane marker, mScarletI-AtLTI6b is detected at the division plane after Venus-MpROP, suggesting that MpROP could play an active role during cell plate formation (Figure 2H). Taken together, the polar localisation of Venus-MpROP in epidermal cells at the edge of the gemma primordia, and early localisation to the division plane in dividing cells support the hypothesis that MpROP regulates cell growth and division during gemma morphogenesis.

### MpROP controls polarised cell growth during gemma morphogenesis

To determine how MpROP influences cell growth and division during gemma morphogenesis, morphometric analyses were performed on wild-type, Mp*rop-1*, and Mp*rop-2* gemmae. The dimensions of a gemma can be defined by gemma height – the length of the proximodistal axis (the distance from the proximal stalk attachment point to the distal edge of the gemma) – relative to gemma width – the distance between the left and right sides of the gemma defined by a line perpendicular to the proximodistal axis (Figure 3A). Mp*rop-1* gemmae are significantly smaller than wild-type gemmae in terms of height and width (Figure 3A, B, D). This difference is due, in part, to defective cellular organisation in Mp*rop-1* as revealed by optical cross section images of wild-type and Mp*rop-1* gemmae. Wild-type gemmae taper from the centre to the edge forming a flattened organ (Figure 3A). There are multiple cell layers in the centre and a single cell layer at the edge and the gradual change in cell layer number from the centre to edge accounts for the taper. This tapering is not observed in Mp*rop-1* gemmae (Figure 3B). Consequently Mp*rop-1* gemmae lack a clear single cell layered edge and there is no clear boundary between the future dorsal and ventral surfaces.

**Figure 3.**
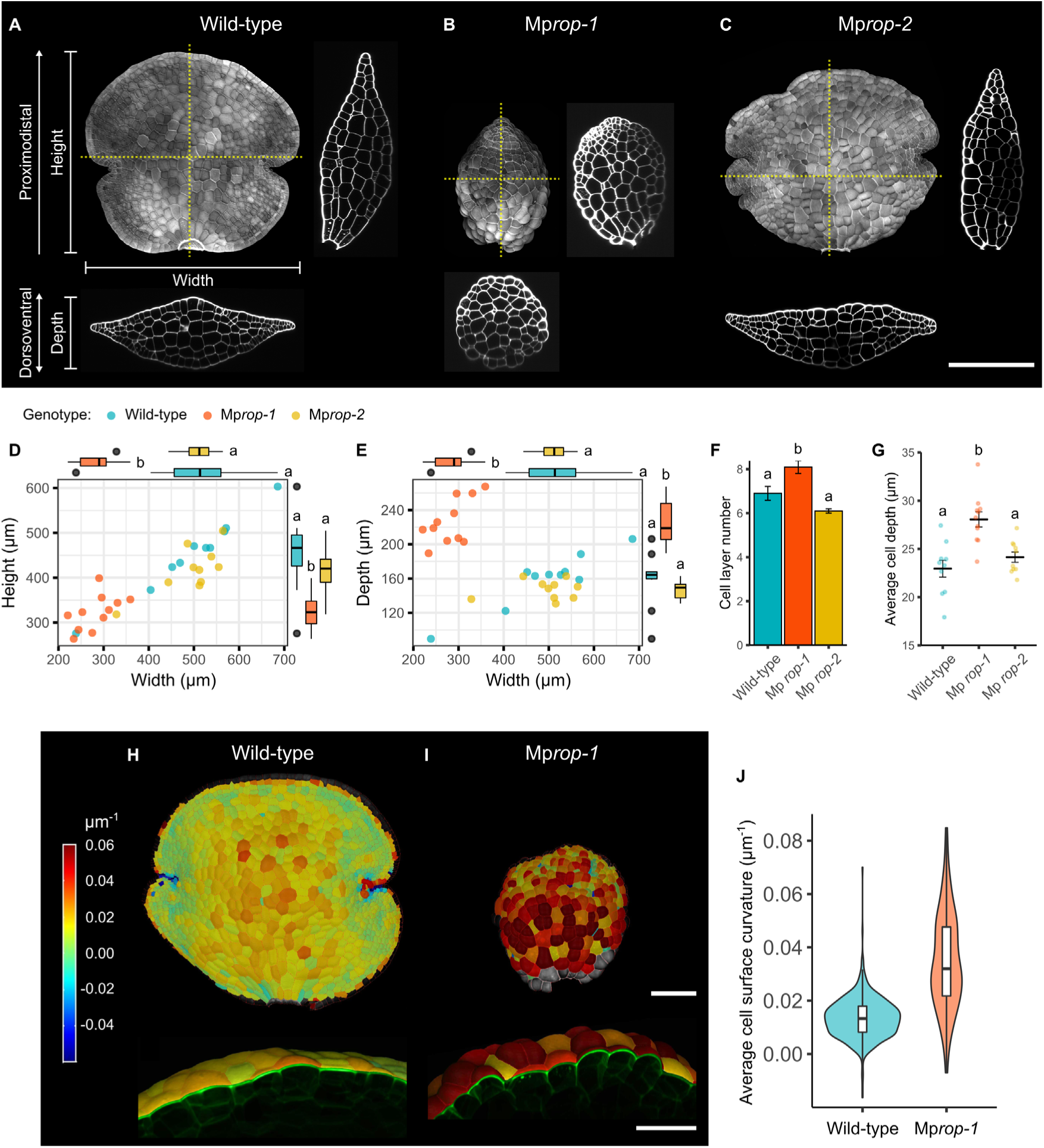
MpROP controls gemma morphogenesis by regulating anisotropic diffuse cell growth. (A-C) Maximum Z-projection image and a pair of orthogonal optical cross section images for (A) wild-type, (B) Mp*rop-1*, and (C) Mp*rop-2* gemmae, which were fixed, cleared, and stained with the cell wall dye SR2200. Yellow dotted lines indicate orthogonal cross section planes. Scale bar, 200μm. (D) Mp*rop-1* gemmae are smaller than wild-type and Mp*rop-2* gemmae in terms of height and width. Each dot in the scatter plot represents height and width of an individual gemma. Box plot above represents distribution of gemma width and box plot to the right represents distribution of gemma height. (E) Mp*rop-1* gemmae are thicker in depth than wild-type and Mp*rop-2* gemmae. Each dot in the scatter plot represents depth and width of an individual gemma. Box plot above represents distribution of gemma width and box plot to the right represents distribution of gemma depth. (F) Mp*rop-1* gemmae consist of more cell layers on average than wild-type and Mp*rop-2* gemmae. By inspecting optical cross section images, the number of cell layers making up the thickest part of each gemma was counted. Bars represent mean and error bars represent standard error of the mean. (G) Mp*rop-1* gemma cells are thicker on average than wild-type and Mp*rop-2* gemma cells. Average cell depth for each gemma was calculated by dividing gemma depth with the number of cell layers at the thickest part of a gemma. Main horizontal black bars represent mean and error bars represent standard error of the mean. Each dot represents average cell depth for an individual gemma. Different lower-case letters (in D-G) indicate groups with significantly different means, based on one-way ANOVA followed by Tukey’s HSD test (p < 0.05). n = 10 for wild-type and Mp*rop-2*, n = 11 for Mp*rop-1*. (H, I) Heatmap displaying average surface curvature for a region 20μm in radius from the centre of each segmented cell surface. Surface view shown on top and optical cross-sectional view on the bottom. Scale bars, 100μm (top row) and 50μm (bottom row). (J) Average cell surface curvature is significantly greater and more varied in Mp*rop-1* than in wild-type. Violin plots show distribution of curvature values and box plots within show median and interquartile range. This analysis was restricted to cells with an area > 150μm^2^ as a region of 20μm in radius greatly exceeds the area of very small cells, meaning curvature values for very small cells represent the tissue curvature of neighbourhood centred around the cell, rather than the surface curvature of the individual cell. Statistically significant difference in mean average cell curvature (Welch’s t-test, p < 2.2e^-16^) and variance (Levene’s test, p < 2.2e^-16^) between wild-type and Mp*rop-1*. n = 523 cells (1 gemma) for wild-type and 199 cells (1 gemma) for Mp*rop-1*.

These data support the hypothesis that MpROP function is required to regulate cell growth and cell division orientations for the formation of the single cell layered gemma edge. The function of MpROP in defining the gemma edge is supported by the observation that Venus-MpROP is polar-localised in cells at the edge of the gemma primordia (Figure 2B-E). Taken together, these data suggest that the polar localisation of MpROP protein at the outer face of cells at the gemma primordia edge is required for the morphogenesis of this flattened structure with tapered edges, characteristic of wild-type gemmae.

The dimensions of the dorsoventral axis can be defined by the gemma depth (thickness) – the distance between the two flattened surfaces at the thickest point, orthogonal to the proximodistal axis (Figure 3A). Although smaller in height and width, Mp*rop-1* gemmae are significantly thicker than wild-type gemmae (Figure 3E). This increased gemma thickness could be due to an increase in cell layer number and, or cell depth compared to wild-type. To test which of these are responsible for increased gemma thickness in Mp*rop-1*, cell layer number and average cell depth (gemma thickness/cell layer number) were quantified. There are more cell layers in Mp*rop-1* gemmae than in wild-type gemmae (Figure 3F). However, the Mp*rop-*1 average cell depth is also greater than wild-type (Figure 3G). Therefore, an increase in both cell layer number and cell depth contributes to thicker gemmae in Mp*rop-1*, consistent with our hypothesis that MpROP regulates both cell growth and cell division orientations during gemma morphogenesis. These parameters were indistinguishable between Mp*rop-2* and wild-type through the morphometric analyses, indicating that the partial loss of function mutant broadly retains the ability to control gemma morphogenesis. The morphometric comparison of wild-type and Mp*rop-1* gemma phenotypes supports our hypothesis that MpROP coordinates organogenesis by regulating both cell growth and cell division orientation.

To test the hypothesis that MpROP controls organogenesis by regulating cell growth orientation, we defined the role of MpROP in shaping gemma epidermal cells. Compared to the smooth future dorsal and ventral surfaces of wild-type gemmae, the overall surface of Mp*rop-1* mutant gemmae is uneven because the epidermal cells swell out of the epidermal surface (Figure 3A, B). To compare the extent of cell growth at the epidermal surface in wild-type and Mp*rop-1*, we estimated the surface curvature of individual gemma epidermal cells using the MorphoGraphX software^29^. The average surface curvature of Mp*rop-1* gemma epidermal cells is significantly greater than that of wild-type, indicating that epidermal cell growth in Mp*rop-1* is not restricted to the plane parallel to the future dorsal and ventral surfaces, but occurs isotropically (Figure 3H-J). Moreover, a wider range of cell surface curvature values are observed for Mp*rop-1* gemma than for wild type, which is consistent with a loss in regulation of cell growth orientation (Figure 3J). Therefore, we conclude that MpROP contributes to gemma morphogenesis by regulating the orientation of anisotropic diffuse cell growth.

### Overexpression of MpROP causes cell division defects during gemma morphogenesis

The defects in cell division orientation in the Mp*rop-1* mutant gemmae are consistent with the hypothesis that MpROP is required to orient the cell plate during cell division. However, the defective orientation of the cell wall in Mp*rop-1* could be a secondary consequence of deregulated cell growth orientation in the loss of function mutant. To determine if ROP signalling directly regulates cell division during gemma development, we disrupted ROP polarity by overexpressing *Venus-*Mp*ROP* under the constitutive promoter, *_pro_*Mp*EF1α*, and studied its effect on gemma morphogenesis.

The characteristic flattened structure of wild-type gemmae is generally maintained in gemmae that overexpress Venus-MpROP. However, the gemma edge is uneven compared to that of wild-type gemmae which is smooth (Figure 4A, E). Unlike the edge of wild-type gemmae, which comprises a single cell layer (Figure 4B), the edge of the *_pro_*Mp*EF1α*: *Venus-*Mp*ROP* gemmae comprises multiple cell layers (Figure 4F). This indicates that overexpression of MpROP causes defective cell division orientation. At the gemma epidermis, cell division orientation is predominantly anticlinal (where the new cell wall is oriented perpendicular to the local organ surface) in wild-type (Figure 4C), however, in *_pro_*Mp*EF1α*: *Venus-*Mp*ROP*, there is clear evidence of periclinal cell divisions (where the new cell wall is oriented parallel to the local organ surface), resulting in cells protruding out of the plane of the epidermal surface (Figure 4G). Moreover, cell wall stubs – where a complete cell wall has not been formed at cytokinesis – are present in *_pro_*Mp*EF1α*: *Venus-*Mp*ROP* gemmae (Figure 4F-H). This demonstrates the disruption of MpROP signalling directly impairs cytokinesis. The cell division defects in gemmae overexpressing Venus-MpROP therefore suggest that the spatiotemporal regulation of MpROP function is necessary for orienting and forming the cell plate during cell divisions which take place during gemma morphogenesis.

**Figure 4.**
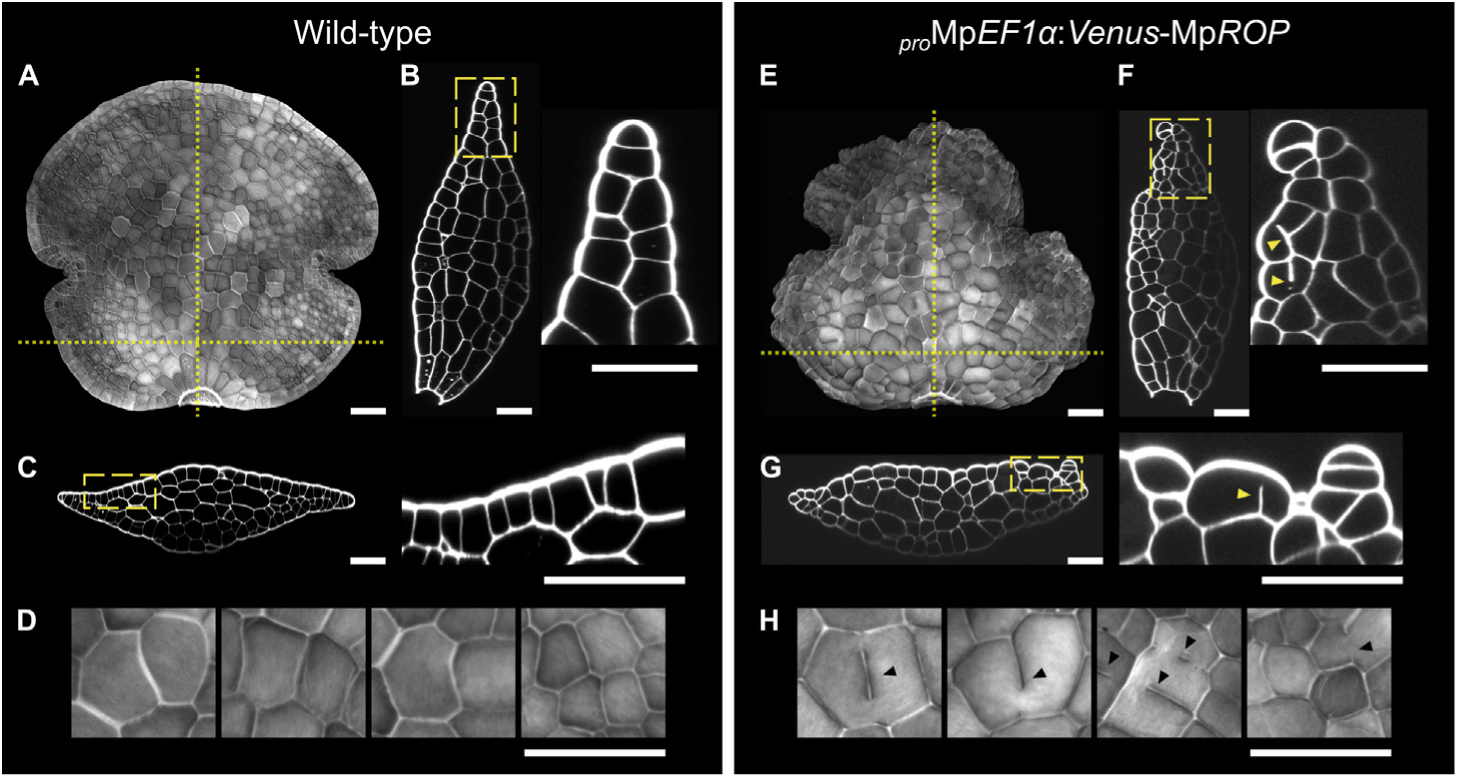
Overexpression of Mp*ROP* causes cell division defects during gemma morphogenesis. (A, E) Maximum projection images of wild-type and *_pro_*Mp*EF1α*: *Venus-*Mp*ROP* gemmae, fixed, cleared, and stained with the cell wall dye SR2200. Dotted yellow lines mark the optical cross section planes of images in B, C, F, and G. (B, F) Optical cross section images (in the YZ plane) of wild-type and *_pro_*Mp*EF1α*: *Venus-*Mp*ROP* gemmae in A and E. Magnified images of region enclosed in dashed yellow line provided. (C, G) Optical cross section (in the XZ plane) images of wild-type and *_pro_*Mp*EF1α*: *Venus-*Mp*ROP* gemmae in A and E. Magnified images of region enclosed in dashed yellow line provided. (D, H) Magnified images of epidermal surface shown in A and E. Cell wall stubs, indicating incomplete cytokinesis was frequently observed in *_pro_*Mp*EF1α*: *Venus-*Mp*ROP* gemmae but never in wild-type gemmae. Arrowheads in F-H mark cell wall stubs. All scale bars, 50μm.

### Venus-MpROP localisation patterns during air chamber development suggest MpROP coordinates tissue development through regulating polarised cell growth and cell division

To investigate how ROP contributes to the formation of complex tissue, we examined MpROP function during air chamber morphogenesis. Air chambers are intricate multicellular structures, which make up the dorsal side of the *M. polymorpha* thallus. A series of stereotypic oriented divisions in the epidermal and the subepidermal cell layers takes place during air chamber formation^30^. We therefore predicted that a defect in cell division pattern causes the formation of defective air chambers in Mp*rop* mutants (Figure 1J-L). To investigate the role of MpROP in air chamber development, we first tested the hypothesis that Venus-MpROP would mark the sites of polarised growth and cell division during air chamber development in wild-type.

The earliest sign of air chamber morphogenesis is the formation of an initial pit (depression region) in the epidermis at the intersect of usually four (sometime 3-5) epidermal cells, close to the apical cell within the meristematic notch (Figure 5A, arrowhead). From this pit, schizogenous separation of the anticlinal walls of neighbouring epidermal cells takes place, giving rise to the initial aperture (Figure 5B, arrowhead). Before schizogenous cell wall separation, Venus-MpROP is localised to the outer epidermal surface, suggesting MpROP promotes polarised outgrowth to create the initial pit (Figure 5G, arrow pointing up). Venus-MpROP also marks the plane where the new inner periclinal wall is expected to form, which will separate the epidermal and sub-epidermal cell layers, consistent with our hypothesis of MpROP involvement in cell division (Figure 5G, arrowhead). After schizogenous cell wall separation, cells which surround the initial aperture then grow in a polarised fashion, inwards and slightly upwards, effectively occluding the aperture at the epidermal surface (Figure 5C). In these aperture-surrounding cells, Venus-MpROP is polarised to the cortical sites of growth, suggesting that MpROP promotes local polarised growth (Figure 5H, arrows). Subsequently, these aperture-surrounding cells divide obliquely to form the roof mother cells, which in turn divide anticlinally to form the air chamber roof cells (Figure 5C, D). Venus-MpROP marks the planes of the oblique division as well as the following anticlinal divisions (Figure 5I, J arrowheads). These data are consistent with the hypothesis that MpROP is required for coordinating the orientations of cell growth and cell division for air chamber roof formation.

**Figure 5.**
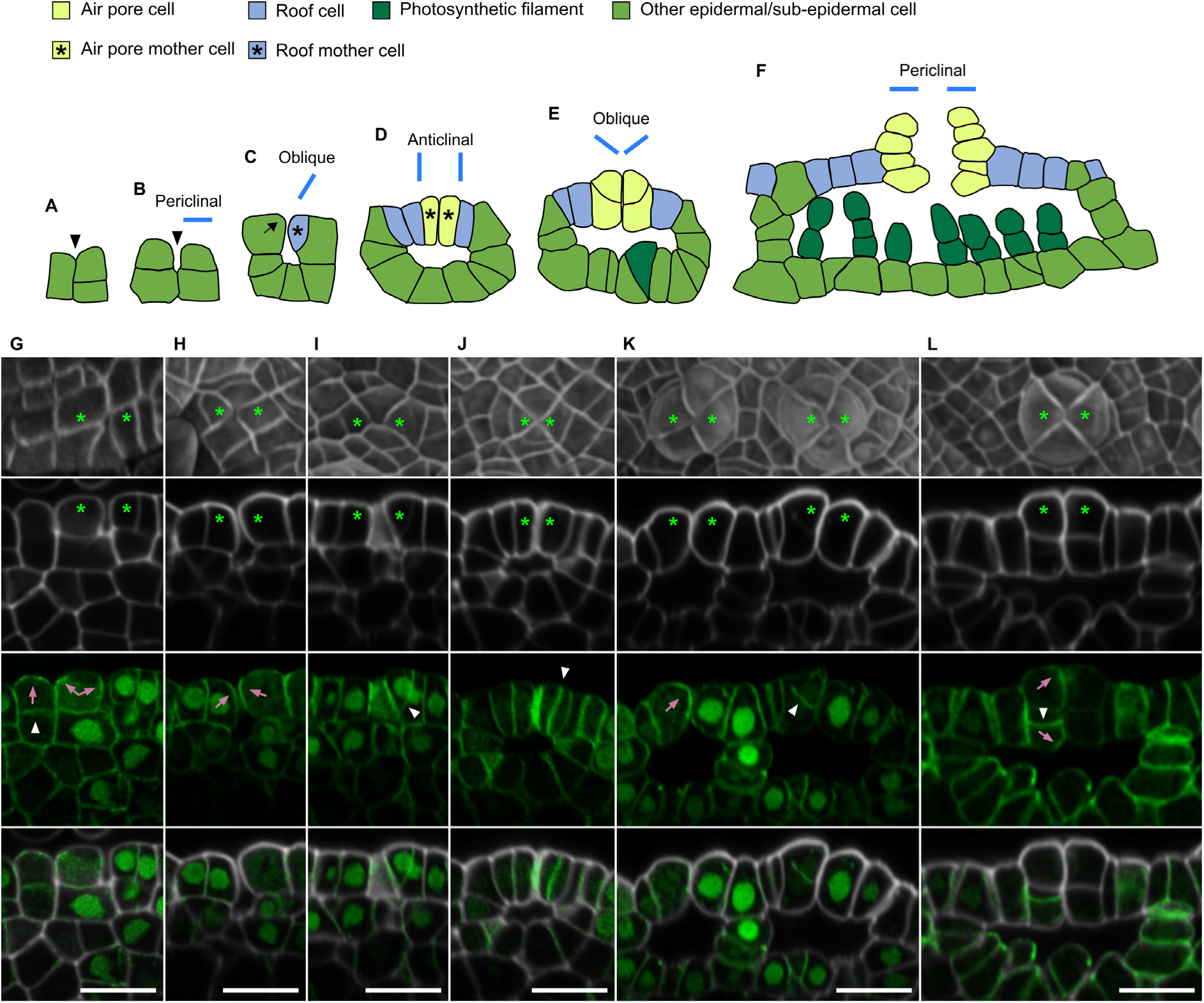
Polar localisation of Venus-MpROP in cells during air chamber formation. (A-F) Schematic representation of wild-type air chamber development in *Marchantia*, based on Apostolakos et al., 1982 and our own observations. Diagrams illustrate how air chambers appear in the cross-sectional plane, perpendicular to the dorsal epidermal surface. (A) Air chamber development commences with the formation of the initial pit (arrowhead) at the intersect of usually four epidermal cells close to the apical cell within the meristematic notch. (B) Schizogenous separation of the anticlinal cell walls at the junction between the four cells results in initial aperture (arrowhead) formation. (C) The epidermal cells surrounding the aperture entrance grow inwards and slightly upwards towards the aperture entrance at the epidermal surface. Arrow indicates cell growth orientation. This results in the closure of the aperture entrance. An oblique division follows producing the roof mother cell. (D) The air chamber roof is formed through the anticlinal division of the roof mother cells. Simultaneously, the anticlinal division of the subepidermal cells which form the air chamber floor, expands the air chamber. Roof cells surrounding the aperture entrance elongate perpendicular to the epidermal surface to differentiate into air pore mother cells. (E) Air pore formation initiates with the oblique division of the air pore mother cells. Development of photosynthetic filaments start at around the same time. (F) Periclinal divisions complete air pore formation. (G-L) Immature air chambers at different developmental stages from 6-day old gemmaling expressing *_pro_*Mp*ROP:Venus-*Mp*ROP* fixed, cleared, and stained with the cell wall dye SR2200. Top row displays surface projection images (SR2200 fluorescence in grey). Second to forth rows show optical cross section images of SR2200 fluorescence (grey), Venus fluorescence (green) and the two channels merged. Asterisks mark cells surrounding the air chamber aperture. Pink arrows mark Venus-MpROP localisation to expected sites of polarised growth. White arrowheads mark Venus-MpROP localisation to newly forming cell plate (indicated by weak/no cell wall staining). All scale bars, 20μm. All images processed in MorphoGraphX.

Air pore formation is initiated by the oblique division of the air pore mother cells, which are the epidermal roof cells which immediately surround the closed aperture (Figure 5E). The oblique division is preceded by a change in cell growth orientation of the air pore mother cells. Venus-MpROP is polar localised to the upper inner (aperture facing) corner of the air pore mother cell, where polarised growth is expected to take place (Figure 5K, arrow). Furthermore, Venus-MpROP marks the newly forming cell plate during the formative oblique division (Figure 5K, arrowhead). At a later stage in air pore formation, Venus-MpROP continues to mark the expected sites of polarised growth and the periclinal division plane of the air pore cells (Figure 5L). Taken together, the Venus-MpROP subcellular localisation pattern at various stages of air chamber development suggests MpROP coordinates this intricate morphogenetic process by regulating local polarised cell growth and cell division.

### MpROP is required for controlling cell division orientation during air chamber development

To test if MpROP regulates cell division pattern during air chamber development, we defined cellular organisation of air chambers in wild-type, Mp*rop-1*, and Mp*rop-2*. The epidermal surface of a mature wild-type air chamber comprises a single layer of roof cells, centred around a tier of air pore cells circumferentially arranged around a narrow aperture (Figure 5F, 6A, D). Neighbouring air chamber units are separated by a wall of cells which stretches from the epidermal surface to the floor of the air chambers. In Mp*rop-1* a wall of cells which form the boundary between neighbouring air chambers are present, however, neither roof cells nor air pore cells were found, confirming that a complete loss of MpROP function prevents the differentiation of these cell types (Figure 6B, E). In wild-type, an oblique division of epidermal cells surrounding the initial aperture results in the differentiation of roof mother cells, which in turn give rise to roof cells (Figure 5C). No evidence of the oblique formative division for roof mother cell differentiation was observed in Mp*rop-1*, suggesting that MpROP is required for the oblique division to initiate air chamber roof formation (Figure 6E).

**Figure 6.**
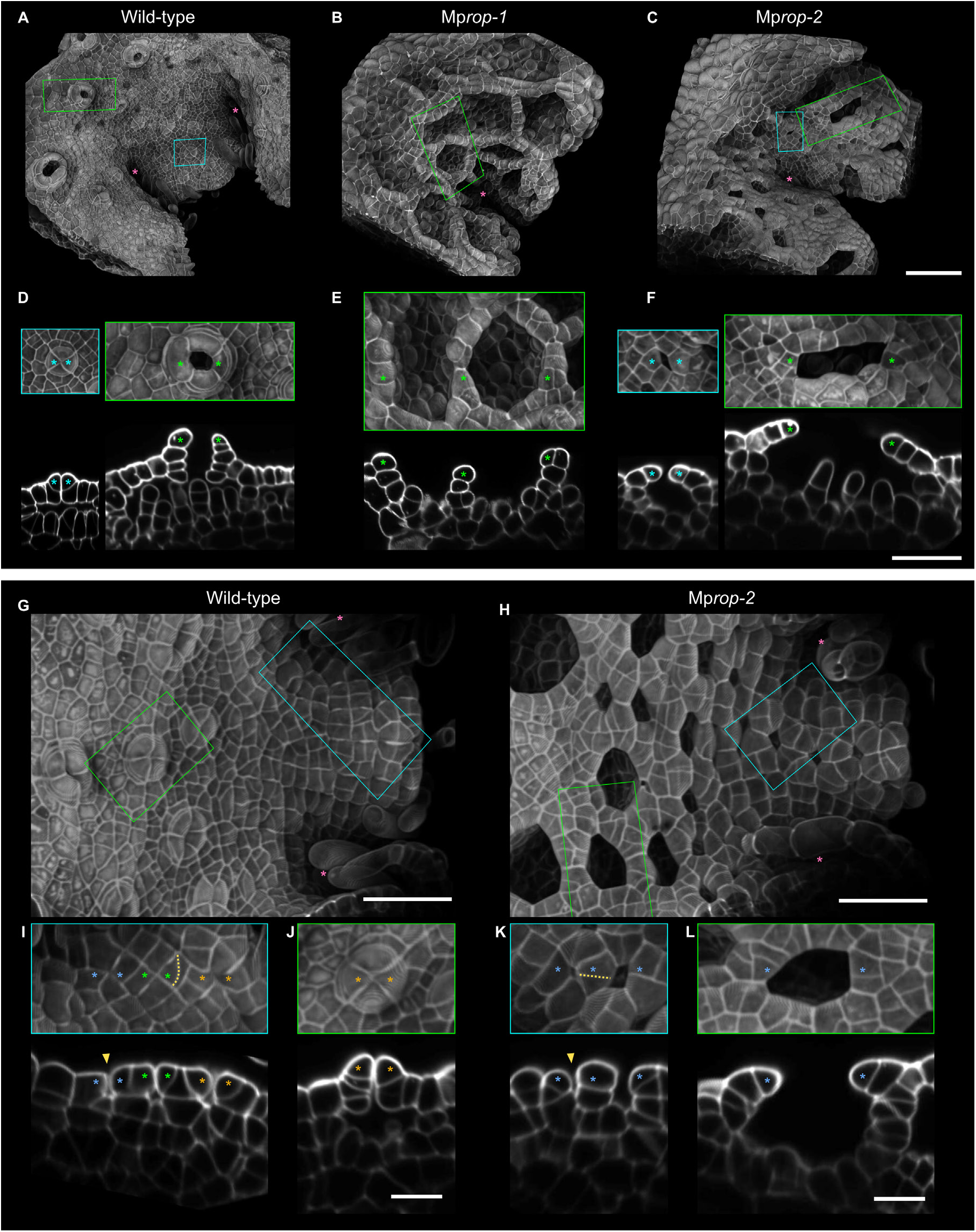
Mp*ROP* is required for switch in cell division orientation during air chamber morphogenesis. (A-C) 3D surface projection images of the dorsal thallus epidermis close to the meristematic notch(es) from 6-day old gemmalings of wild-type, Mp*rop-1*, and Mp*rop-2* (fixed, cleared, and stained with the cell wall dye SR2200). Higher magnification images of boxed regions are show in D-F. Pink asterisks mark meristematic notches. (D-F) Surface projection (top row) and optical cross section (bottom row) images of immature and mature air chambers from wild-type, Mp*rop-1*, and Mp*rop-2*. In wild-type, anticlinal cell divisions form the air chamber roof. This takes place in Mp*rop-2* but not in Mp*rop-1*. In wild-type, a change in cell division orientation from anticlinal to periclinal facilitates air pore formation. This was not observed in Mp*rop-1* or Mp*rop-2*. (G, H) 3D surface projection images of a relatively flat region of dorsal thallus epidermis, sandwiched between two meristematic notches (marked with pink asterisks), which bifurcated recently from 5-day old gemmalings of wild-type and Mp*rop-2*. Higher magnification images of boxed regions are show in I-L. (I-L) Surface projection (top row) and optical cross section (bottom row) images of regions marked in G and H. (I, K) Close to the meristematic notches, an initial aperture (yellow arrowhead) forms at the intersect of ∼4 epidermal cells. Polarised growth of aperture surrounding cells occlude the initial aperture in wild-type. Polarised growth fails to take place in Mp*rop-2* and aperture remains open. In wild-type, the division of aperture surrounding cells is oriented radially with respect to the closed aperture (yellow dotted line). In Mp*rop-2*, division of aperture surrounding cells can be misoriented by 90° (yellow dotted line). (J, L) Further from the meristematic notches (i.e. later in development), air chamber apertures remain closed during air pore formation in wild-type. In Mp*rop-2*, air chamber apertures have enlarged and the number of aperture surrounding cells have increased. In A-C and G-H, pink asterisk mark the meristematic notch, whilst in D-F and I-L, asterisks mark epidermal cells forming the aperture boundary visible in the optical cross section images. Scale bars, 100μm (A-C), 50μm (D-H), 20μm (I-L). All images processed in MorphoGraphX.

Cellular organisation in the air chambers of the Mp*rop-2* mutant also indicates defects in cell divisions. Before air pore formation starts in wild-type, the roof cells surrounding the occluded aperture elongate perpendicular to the epidermal surface and differentiate into the air pore mother cells (Figure 5D, K). These then undergo an oblique division to produce the first air pore cells, which continue to divide periclinally, to form the final air pore (Figure 5E, F, 6D). Although there is evidence of anticlinal divisions of roof cells in Mp*rop-2*, neither oblique nor periclinal divisions of roof cells surrounding the aperture were observed (Figure 6F). Moreover, Mp*rop-2* roof cells did not elongate perpendicular to the epidermal surface, as observed in wild-type, and consequently these cells did not differentiate into air pore mother cells. Therefore, MpROP activity is likely also required for the oblique divisions that initiate air pore formation.

As well as lacking the multicellular air pore, which in wild-type encircles the aperture, the apertures of Mp*rop-2* air chambers are much larger than that of wild-type (Figure 1I, K, 6A, C). To determine the cause of the enlarged air chamber apertures of Mp*rop-2*, we compared wild-type and Mp*rop-2* development just after the formation of the initial aperture. Following schizogenous cell wall separation which forms the initial aperture, the aperture-surrounding cells in wild-type grow in a polarised fashion to occlude the initial aperture (Figure 6I). By contrast, polarised growth of aperture-surrounding cells is less evident in Mp*rop-2*, resulting in the aperture staying open (Figure 6K). This suggests MpROP is required for the local polarised growth of aperture-surrounding cells and is consistent with the polar localisation of Venus-MpROP in equivalent aperture-surrounding cells in wild-type (Figure 5H). After the closure of the initial aperture in wild-type, the aperture reopens only at the final stages of air chamber morphogenesis, when air pore formation is being completed (Figure 6A, G). By contrast, the Mp*rop-2* air chamber aperture increases in size during air chamber development (Figure 6C, H). The wild-type aperture remains closed presumably because growth of aperture surrounding cells, not only during the initial closure of the aperture but also later during roof and pore formation, is polarised and oriented towards the closed aperture. Enrichment of Venus-MpROP signal at the aperture facing surfaces of aperture surrounding cells throughout wild-type air chamber development is consistent with a role of MpROP in promoting polarised growth towards the aperture (Figure 5). When these aperture surrounding cells divide, the new cross wall is placed perpendicular to the cell growth axis, resulting in only one of the daughter cells facing into the aperture (Figure 6G, I, J). When the dorsal epidermal surface is viewed from above, the newly placed cross wall appears to bisect the epidermal surface of the parental cell, radially with respect to the closed aperture (Figure 6I, yellow dotted line). Therefore, by orienting polarised growth and cell division towards the aperture, the aperture remains closed and the number of cells surrounding the aperture remains constant (usually 4). In Mp*rop-2*, division of some aperture surrounding cells are clearly misoriented with respect to the aperture (Figure 6H, K, L). New cross walls are seen bisecting the parental wall which faces into the aperture (in other words, division misoriented by 90°, Figure 6K, yellow dotted line). These misoriented divisions of aperture surrounding cells can be observed shortly after the formation of the initial aperture and continues to be observed during air chamber development in Mp*rop-2*. Therefore, unlike wild-type where the number of cells forming the aperture perimeter remain constant, the number of cells forming the aperture perimeter in Mp*rop-2* increases during air chamber development. This likely contributes to the formation of larger apertures in Mp*rop-2* than in wild-type. Thus, defects in polarised cell growth and cell division orientation causes enlarged apertures in Mp*rop-2* from an early stage in air chamber morphogenesis which become progressively larger.

To summarise, based on Venus-MpROP polar localisation to sites of growth and division in wild-type, combined with defective polarised growth and cell division orientation in Mp*rop* mutants, we conclude MpROP likely controls the subtle changes in cell shape and intricate switches in cell division orientation during the morphogenesis of the air chambers.

## DISCUSSION

### ROP protein is required for orchestrating morphogenesis of complex plant tissues and organs

We report that ROP function is required for the morphogenesis of complex tissues and organs in the liverwort *M. polymorpha*. During air chamber and gemma morphogenesis, Venus-MpROP marks the sites of local polarised growth at the cell surface and the expanding cell plate of cells undergoing cytokinesis in wild-type. This is consistent with the defects in polarised cell growth and cell division orientation in loss of function Mp*rop* mutants. Our data demonstrate that the regulation of local polarised cell growth and cell division orientations by the ROP protein is critical for the formation of complex plant tissues and organs.

### The single Mp*ROP* gene is required for both tip growth and anisotropic diffuse growth

By correlating the polar localisation patterns of Venus-MpROP with the cell shape defects in Mp*rop* loss of function mutants, we demonstrated that ROP function is required for both tip growth and anisotropic diffuse growth in *M. polymorpha*. We show that ROP function in regulating anisotropic growth is important for tissue development and organogenesis. Although ROP function in tip growth has been previously described in both vascular plants^4,5,12^ and bryophytes^14,23,31^, ROP function in anisotropic diffuse growth had only been described in *A. thaliana*^7,8^. To our knowledge, our study is the first to clearly demonstrate ROP function in anisotropic diffuse growth in any bryophytes. Our findings suggest that ROP function in regulating anisotropic diffuse growth is conserved within land plants. ROP proteins in *A. thaliana* direct anisotropic diffuse growth by governing the organisation of the actin and microtubule cytoskeleton^7,8,32^. It was recently shown that the anisotropic organisation of cortical microtubules is disrupted in epidermal cells of a Mp*rop* loss of function mutant^23^. Moreover, the Mp*rop-1* mutant gemma epidermal cells swelling out of the epidermal surface (Figure 3B, I) resemble *A. thaliana* sepal epidermal cells treated with the microtubule depolymerising drug oryzalin^33^. It is therefore likely that MpROP also regulates anisotropic diffuse growth by coordinating the organisation of the cytoskeleton, through a mechanism broadly conserved in land plants.

By demonstrating the requirement of the single *M. polymorpha ROP* gene in regulating both tip growth and anisotropic diffuse growth, we have shown that a single *ROP* gene is sufficient for different forms of polarised cell growth. In *A. thaliana*, which encodes 11 *ROP* genes in its genome, distinct ROP proteins, localised differentially within a cell, carry out distinct functions to regulate polarised cell growth. For example, in pavement cells, AtROP2 and AtROP4 promote lobe outgrowth whilst AtROP6 functions antagonistically, restricting outgrowth at the neck region, contributing to formation of the jigsaw puzzle shaped cells^7,8^. And in root hair cells, AtROP2 localises to the apex to promote tip growth^4^, whilst AtROP10 localises to the shank to regulate cell wall deposition for shank hardening^34^. It is unclear how seemingly analogous processes like root hair and rhizoid tip growth, require multiple types of ROP proteins in *A. thaliana* but only a single type of ROP protein in *M. polymorpha*. Undoubtedly, a range of ROP regulators and effectors is critical for differential regulation of ROP signalling in distinct cell types to achieve different forms of polarised cell growth. The ability of ROP regulators to control very specific aspects of ROP function is exemplified by MpRopGEF, a positive regulator of MpROP, which is specifically required to initiate gemma development^35^. There are, however, likely limits to the capability of a single *ROP* gene. For instance, with just a single *ROP* gene, it is implausible that distinct, antagonistically functioning ROP domains could be generated within the same cell, as seen in the *A. thaliana* pavement cells. This may explain why cells with multiple sites of polarised growth like pavement cells and trichomes are absent in *M. polymorpha*. Therefore, a single *ROP* gene is sufficient for regulating different forms of polarised cell growth, however, we predict that neofunctionalisation, prompted by the expansion of the *ROP* gene family in some plant lineages, resulted in greater spatial control over polarised cell growth and facilitated elaboration of cellular morphogenesis.

### ROP regulates tissue development and organogenesis by controlling cell division orientation

We demonstrate that ROP protein regulates cell division orientation, and that this function of ROP is critical for tissue morphogenesis and organogenesis. In Mp*rop* loss of function mutants, formative oblique (asymmetric) cell divisions during air chamber development are defective, preventing the formation of the air chamber roof and the multicellular air pore. Previous reports of cell division defects in *rop* loss of function mutants in other species have focussed on asymmetric cell divisions and it was unclear if ROP regulates orientation of specific division events or it has a more general role in regulating a wide range of division events^9,10^. As well as defects in oblique divisions, we provide evidence of defects in the orientation of symmetric cell divisions in Mp*rop* loss of function mutants, for example during gemma morphogenesis in Mp*rop-1* and air chamber roof development in Mp*rop-2*. Therefore, ROP function is required for orienting both symmetric and asymmetric divisions during plant morphogenesis.

How precisely ROP regulates cell division orientation remains unclear. We predict ROP influences cell division orientation in several ways. First, ROP proteins could be influencing cell division orientation indirectly via regulating cell growth orientation. Asymmetric divisions for initiating *M. polymorpha* air chamber roof formation as well as *P. patens* protonema branching are proceeded by polarised cell growth and the new cell plate is placed perpendicular to the cell growth axis^10^. Defects in the orientation of these divisions in the Mp*rop-2* and the Pp*rop2 3 4* mutants are both accompanied by defective polarised growth. Furthermore, on occasions when Pp*rop2 3 4* manages to initiate polarised outgrowth for branching, division orientation is oblique, suggesting that defective cell division orientation is, at least partially, a consequence of defective polarised growth^10^. Second, ROP could be determining division plane position by controlling nuclear position. For asymmetric divisions in both *Z. mays* and *P. patens*, ROP has been suggested to direct actin mediated nuclear migration to asymmetrically position the nucleus before division^9,10^. Third, ROP could be involved in determining the cortical division zone in a more direct manner. Together with the mitosis specific kinesin, POK, REN proteins which negatively regulate ROP signalling, have been proposed to act just before and/or during division to determine division plane orientation^28,36^. How POK and REN function in division plane selection is not well understood, however, it implies spatially confined deactivation of ROP just before or during division is necessary for correct division plane selection. This is consistent with defects in division orientation we observed in *_pro_*Mp*EF1α*: *Venus-*Mp*ROP* gemmae, where MpROP is overexpressed. Finally, some of the cell division orientation defects and the tissue defects of Mp*rop* mutants are likely to be interdependent. Defects in cell division orientation results in defective cellular organisation. This could influence the distribution of mechanical tension within the tissue which in turn could influence cell division orientation^37^. In wild-type *M. polymorpha*, the air pore – the multicellular structure comprising a tier of cells circumferentially arranged around a narrow aperture – likely acts as a load bearing structure and we predict the absence of air pores in Mp*rop* mutants alters the mechanical stress patters within the tissue. The defectively oriented cell divisions which result in enlarged air chamber apertures in Mp*rop-2*, could be caused by alterations in tissue mechanics compared to wild-type. Taken together, we predict ROP influences cell division orientation in multiple ways.

### ROP as a coordinator of polar processes

We demonstrated MpROP function in polarising cell growth and orienting cell divisions, mainly through characterising loss of function Mp*rop* mutants. Disrupting ROP polarity through overexpressing Mp*ROP*, caused the formation of cell wall stubs, implying that proper spatial regulation of ROP is necessary not only for orienting but also for constructing the cell plate. Polarised cell growth and cytokinesis in plants are both polar processes dependent on polarised vesicle trafficking^36,38^. We speculate that by polar localising to different cellular compartments during the cell cycle, ROP orchestrates cytoskeletal organisation to direct exocytosis to a cortical site, during polarised growth, and cytokinetic vesicles to the division plane, during cell plate expansion, thereby ensuring proper spatiotemporal coordination of cell growth and division. We propose this ability to regulate polarised cell growth and cell division pattern in a coordinated manner makes ROP instrumental for morphogenesis of complex plant structures.

## MATERIALS AND METHODS

### Plant material and growth conditions

Transgenes were introduced into sporelings generated from a cross between the Takaragaike-1 (Tak-1, male) and Takaragaike-2 (Tak-2, female) wild-type accessions of *Marchantia polymorpha*. For phenotypic comparisons with T_1_ mutant plants, wild-type siblings of the T_1_ mutants were used as wild-type (see Figure S3B for detail). Unless stated otherwise, plants were grown on sterile plates containing modified Johnson’s medium 6mM KNO_3_, 500μM MgSO_4_.7H_2_O, 4mM Ca(NO_3_)_2_.4H_2_O, 25μM KCl, 12.5μM H_3_BO_3_, 1μM MnSO_4_.4H_2_O, 1μM ZnSO_4_.7H_2_O, 0.25μM CuSO_4_.5H_2_O, 0.25μM (NH_4_)_6_Mo_7_O_24_.4H_2_O, 25μM FeSO_4_.7H_2_O, 25.5μM FeNaEDTA, 400μM (NH_4_)_2_SO_4_, 600μM NH_4_H_2_PO_4_, 555μM myo-Inositol, 0.5g/L MES hydrate, 1% sucrose, pH adjusted to 5.6) or ½-strength B5 Gamborg’s medium (1.5g/L B5 Gamborg’s, 0.5g/L MES hydrate, 1% sucrose, pH adjusted to 5.5), solidified with 1% plant agar. Plants on agar plates were grown at 23°C under continuous white light (50–60µmol m^-2^ s^-1^).

For crossing, 2–3 weeks old gemmalings grown under the above condition, were transplanted onto autoclaved soil (1:3 mixture of fine vermiculite and John Innes No.2 or Neuhaus N3 compost) in SacO2 Microbox containers. Plants on soil were grown at 20°C under white light (50–60µmol m^-2^ s^-1^) supplemented with far-red light (30–40µmol m^-2^ s^-1^), in long day conditions (16 hours light, 8 hours dark).

### Plasmid construction

#### CRISPR/Cas9 constructs for Mp*ROP* mutagenesis

Within the Mp*ROP* (Mp7g17540) genomic sequence, three target sites, which satisfy the sequence criteria (5’ GN_19_NGG 3’), were selected. BLAST search was performed against the *M. polymorpha* genome using the three target sequences as query, to confirm that these sequences are specific to Mp*ROP*. For each target site, a forward (CTCG N_19_) and a reverse (AAAC rev.comp.N_19_) oligonucleotide was designed (Table S2). To generate entry clones with a single gRNA sequence, the annealed oligonucleotides were ligated with *Bsa*I digested pMpGE_En03^39^. To generate CRISPR expression constructs, LR reaction was performed between the entry clones containing specific sgRNA sequences and the pMpGE010 destination vector which contains the *_pro_*Mp*EF1α:Cas9* cassette^39^.

#### *_pro_*Mp*ROP*:*NLS-Venus*, *_pro_*Mp*ROP*:*Venus-*Mp*ROP* and *_pro_*Mp*EF1α*:*Venus-*Mp*ROP*

The Mp*ROP* promoter (4900bp region upstream of Mp*ROP* start codon), the Mp*EF1α* promoter^41^ (1735bp), and the Mp*ROP* gene (3069bp, from start codon to end of 3’ UTR) sequences were amplified from the genomic DNA of Tak-1. Sanger sequencing confirmed that Mp*ROP* sequences are identical in Tak-1 and Tak-2. The Venus-YFP coding sequence with and without upstream nuclear localisation signal, and the NOS terminator sequence were amplified from constructs kindly provided by Anna Thamm^42^. Phusion High-Fidelity DNA Polymerase (Thermo) or CloneAmp HiFi PCR Premix (Takara) was used to PCR amplify all the above sequences with primers listed in Table S2. In-Fusion cloning (Toyobo) was used to introduce the above amplified sequences into the multiple cloning site of pCAMBIA1300. To generate *_pro_*Mp*ROP*:*NLS-Venus*, the NOS terminator was first cloned into pCAMBIA1300 digested with *Sac*I and *Eco*RI. The resulting construct was digested with *Sal*I and *Sac*I to introduce *_pro_*Mp*ROP* and *NLS-Venus* sequences. To generate *_pro_*Mp*ROP*:*Venus-*Mp*ROP* and *_pro_*Mp*EF1α*:*Venus-*Mp*ROP*, the Mp*ROP* and NOS terminator sequences were first cloned into pCAMBIA1300 digested with *Sac*I and *Eco*RI. The resulting construct was digested with *Sal*I and *Bsr*GI to introduce the *Venus* sequence, together with either the *_pro_*Mp*ROP* or the *_pro_*Mp*EF1α* sequences.

#### *_pro_*Mp*ROP*:Mp*ROP*

The OpenPlant Loop assembly toolkit and associated protocol provided by Sauret-Güeto et al., 2020 were used to generate *_pro_*Mp*ROP*:Mp*ROP* construct for complementation. The Mp*ROP* promoter sequence, corresponding to 4kb directly upstream of the Mp*ROP* start codon was PCR amplified from the *_pro_*Mp*ROP:Venus-*Mp*ROP* construct in four separate fragments to domesticate internal *Bsa*I and *Sap*I recognition sites. Thermo Scientific Phusion Green High-Fidelity DNA Polymerase and primers listed in Table S2 were used. The four PCR products were assembled into the pUAP4 vector through a *Sap*I assembly reaction, to generate L0_PROM5_MpROP. The CDS for Mp*ROP* was ordered as a gBlocks Gene Fragment from IDT. Appropriate overhang sequences were included in the fragment for *Sap*I assembly into the pUAP4 vector, to generate L0_CDS_MpROP. Three L0 parts (L0_PROM5_MpROP, L0_CDS_MpROP, and L0_3TERM_NOS-35S which encodes the Nos-35S terminator) were assembled into the pCk2 acceptor vector through the *Bsa*I assembly reaction to generate a L1 plasmid encoding the transcriptional unit *_pro_*Mp*ROP:*Mp*ROP-Nos35S*. Finally, the L1 plasmid was combined with OP-062, OP-066, OP-012 and OP-005 (all provided in the OpenPlant Kit) in a L2 *Sap*I assembly reaction to generate an expression vector encoding the hygromycin phosphotransferase gene (for selection of transformants), *_pro_*Mp*ROP:*Mp*ROP* and *_pro_*Mp*UBE2:mTurqoise-N7* (ubiquitous nuclear marker).

### Plant transformation

#### Sporeling transformation to introduce CRISPR constructs and Venus encoding constructs into wild-type

Based on method developed by Ishizaki et al., 2008, and adapted by Honkanen et al., 2016. Intact wild-type (Tak-1 x Tak-2) sporangia were harvested in 1.5ml Eppendorf tubes, which were then sealed with micropore tape and left to dry in a SacO2 Microbox container, 1/5 filled with silica gel. After two weeks of drying, Eppendorf tubes containing sporangia were closed and transferred to the −80°C freezer for storage until transformation. To sterilise, thawed sporangia were crushed in 0.1 % (w/v) sodium dichloroisocyanurate (Sigma) solution and left for 2 minutes. The spore suspension was then spun down at 15,000g for 2 minutes to pellet the spores. The supernatant was removed, and spores were resuspended in sterile water. The spore suspension was added to sterile 125ml Erlenmeyer flasks containing 25ml M51C medium^44,45^, to achieve a spore concentration of roughly 1 sporangium per flask. Spores in sealed sterile flasks were cultured for 7 days at 23°C under continuous white light (50–60µmol m^-2^ s^-1^) with constant agitation (130 rpm).

GV3101 *Agrobacterium tumefaciens* transformed with the desired expression construct was cultured from a single colony in 5ml of M51C medium at 28°C in the dark with constant agitation (180rpm). After 2 days incubation, 2ml of the culture was spun down (3,000g, 10 minutes) and the pellet resuspended in 10ml of M51C containing 100μM acetosyringone. After a further 6 hours of incubation, 1ml of the induced *Agrobacterium* culture was added to the 7-day old sporeling culture, together with acetosyringone (final concentration of 100μM), and left to co-cultivate for 2 days at 23°C under continuous white light (50–60µmol m^-2^ s^-1^) with constant agitation (130 rpm). The liquid co-culture was passed through a 40μM nylon cell strainer (Corning) and captured sporelings were washed with sterile water. Sporelings were then plated on modified Johnson’s agar plates containing 150μg/ml cefotaxime and 10μg/ml hygromycin for selection.

#### Thallus transformation for complementing Mp*rop* mutants

Based on method developed by Kubota et al., 2013. First, plants were grown from gemmae or thallus clippings (as Mp*rop-1* produces very few gemmae) on Gamborg’s medium under standard growth conditions for two weeks. Then, using a sterilised cork borer, disc shaped clippings (4mm diameter) were collected from the thallus, avoiding the meristematic notch. Thallus discs were placed back on original media plates (making sure ventral side facing the agar surface) for 3 days of regeneration under standard growth conditions.

GV3101 *Agrobacterium tumefaciens* transformed with the complementation construct was cultured and induced as described above for sporeling transformation. The induced *Agrobacterium* culture (1ml) was added to a sterile 125ml Erlenmeyer flask containing 50ml M51C medium, together with acetosyringone (final concentration of 100μM) and 40–50 regenerating thallus discs. Flasks were left at 23°C under continuous white light (50–60µmol m^-2^ s^-1^) with constant agitation (130 rpm) for 3-day co-cultivation. Following 3 days of co-cultivation, thallus discs were washed in water then in 1mg/ml Cefotaxime as described by Kubota et al., 2013, and finally plated on standard Gamborg’s medium containing 150μg/ml cefotaxime and 10μg/ml hygromycin and grown under standard condition for selection. Transformants were selected 2–3 weeks later.

### Genotyping CRISPR mutants

To check for CRISPR/Cas-9 induced mutations, Phire Plant Direct PCR kit (Thermo Scientific) was used with primers listed in Table S2 to amplify parts of the Mp*ROP* genomic sequence, directly from thallus tissue of transformants. Gel purified PCR products were sent for Sanger sequencing with primers listed in Table S2.

To check Mp*ROP* transcript sequence of Mp*rop* mutants, total RNA was extracted from 12-day old gemmalings using RNeasy Plant Mini Kit (Qiagen) with DNase-I digestion. From 1μg of total RNA, cDNA was synthesized in a 20μl reaction with ProtoScript II Reverse Transcriptase (NEB) and oligo(DT) (Sigma) in the presence of Murine RNase inhibitor (NEB), following NEB protocol (#M0368). cDNA diluted 10 times in nuclease-free water was used as template to amplify Mp*ROP* coding sequence, with Phire Plant Direct PCR kit (Thermo Scientific) and primers listed in Table S2. Gel purified PCR products were sent for Sanger sequencing with primers listed in Table S2.

To select *Cas9*-free T_1_ Mp*rop* mutants and wild-type siblings, fresh sporangia produced after crossing T_0_ Mp*rop* mutants to wild-type (Tak-1 or Tak-2) were sterilised in 1 % (w/v) sodium dichloroisocyanurate (Sigma) solution for 3 minutes, before washing three times in sterile water. Sporangia were burst in sterile water to release spores, and these were plated on modified Johnson’s agar plates without antibiotics. After three weeks, hygromycin sensitivity, which indicates the lack of the *Cas9* transgene, was tested by taking two thallus clippings from each selected T_1_ plant and replica plating on Johnson’s agar plates with and without hygromycin. The inheritance of the parental Mp*rop* or Mp*ROP* alleles were assessed by Sanger sequencing for hygromycin sensitive T_1_ plants with Mp*rop* or wild-type phenotypes, respectively. For those where the inheritance of the parental allele was confirmed, replica plating on plates with and without hygromycin was repeated to verify the lack of the *Cas9* transgene.

### Rhizoid imaging and length measurement

To prepare plates for the rhizoid growth assay, 50ml of autoclaved molten ½-strength B5 Gamborg’s medium containing 0.8% Phytagel was poured in each square petri dish (120×120×17mm). When the dish is propped up vertically, to have a horizontal Phytagel surface on which gemmae can be placed, Phytagel occupying the top 30mm of the dish was cut out neatly, to leave a phytagel surface approximately 5mm in width (corresponds to the thickness of the poured Phytagel), which is perpendicular to the dish surface. Four to five gemmae were plated along this narrow surface and sealed dishes were left vertically under standard growth conditions. Seven and ten days after plating, rhizoids of individuals gemmalings were imaged using the Leica M165 FC stereomicroscope, equipped with the Leica DFC310 FX camera. Maximum rhizoid length, defined as the vertical distance from the phytagel surface, on which the thallus sits, to the tip of the rhizoid which has grown the furthest away from the phytagel surface, was measured for each gemmaling in Fiji.

### Phenotypic characterization of mutant tissue using stereomicroscopy

Wild-type and mutant tissues were imaged with the Leica M165 FC stereomicroscope, equipped with the Leica DFC310 FX camera, or the Keyence VHX7000 digital microscope equipped with the VH-Z00R/Z00T and VH-ZST lenses and the VHX-7020 camera.

### Tissue fixation and clearing

The fixative, 4% (w/v) paraformaldehyde in 1x PBS, supplemented with 0.1% (v/v) Brij® L23, was freshly prepared as described by Ursache et al., 2018. Gemmae and meristematic notches (dissected from 5-6 day old gemmalings) were suspended in the fixative in Eppendorf tubes and fixed under vacuum (2x 30-minute vacuum treatment). After removing the fixative, samples were washed in 1x PBS twice. Samples in Eppendorf tubes were then suspended in the clearing solution, ClearSeeα^48^, and vacuum (1x 30-minute) was applied to promote infiltration. Samples were left in ClearSeeα at room temperature in the dark for at least a week (replacing ClearSeeα with fresh ClearSeeα every 1– 2 days) before imaging.

### Cell wall staining of live and fixed specimens

Live gemmae were stained in 10μg/ml propidium iodide (PI) for 10 minutes. To image very immature gemmae still attached to the base of the gemma cup, transversely dissected gemma cups were stained. PI was washed off with water before confocal imaging.

Fixed specimens were stained in 0.2% (v/v) SCRI Renaissance2200 (SR2200) in ClearSeeα at room temperature in the dark, overnight. SR2200 was replaced with ClearSeeα a few hours before imaging.

### Epifluorescence microscopy

Venus epifluorescence images were acquired on the Leica MZ16FA stereomicroscope equipped with Leica EL6000 mercury lamp and Leica DFC300 FX camera, through the YFP filter (excitation: 500– 520nm; emission: 540–580nm).

### Confocal microscopy

Confocal live imaging was performed on an upright Zeiss LSM780 equipped with GaAsP detectors. All PI stained gemmae expressing *_pro_*Mp*ROP:NLS-Venus* were imaged with a x20/0.8 NA air objective. Overall images of gemmae expressing *_pro_*Mp*ROP:Venus-*Mp*ROP* and *_pro_*Mp*UBE2:mScarletI-*At*LTI6b* were also acquired with a x20/0.8 NA air objective. Higher magnification 16-bit images for fluorescence quantification in gemma epidermal cells and rhizoids were acquired with a x40/1.1 NA water objective. The following excitation laser wavelength and emission capture bandwidth were used: Venus (ex 514nm, em 518–544nm), mScarletI (ex 561nm, em 571–624nm), PI (ex 561nm, em 580–624nm). Sequential scanning was used to avoid bleed through. All specimens were placed in a chamber setup described by Kirchhelle & Moore, 2017 for live imaging. Rhizoid were imaged after growing 0-day old gemmae in the chamber for 24 hours under standard growth conditions.

Confocal imaging of fixed specimens cleared in ClearSeeα was performed on an inverted Zeiss LSM880 equipped with GaAsP detectors. All images were acquired with a 40x/NA 1.2 objective, using silicon immersion oil, which closely matches the refractive index of ClearSeeα. The tile scan function was used to image large specimens and the resulting images were stitched using the online stitching tool in ZEN Black. The following excitation laser wavelength and emission capture bandwidth were used: SR2200 (ex 405nm, em 420–500nm, based on Tofanelli et al., 2019), Venus (ex 514nm, em 518–560nm). Sequential scanning was used to avoid bleed through when imaging both SR2200 and Venus fluorescence. For quantifying surface curvature in MorphoGraphX, 16-bit images were obtained with voxel size of 0.35×0.35×0.35μm. All fixed and cleared specimens were mounted in ClearSeeα in a Gene Frame (Thermo Scientific) attached to a standard microscope slide.

### Fluorescence intensity quantification

To compare Venus-MpROP and mScarletI-AtLTI6b localisation patterns, Venus and mScarletI fluorescence intensities along a plasma membrane region of interest were measured in Fiji. After background subtraction from the sum projection image (sum of 5 consecutive slices), the segmented line tool (line width adjusted to cover the whole width of the plasma membrane, spline fit) was used to manually trace a line over the plasma membrane. Profile plots of the normalised signal intensities of Venus and mScarletI along the traced line were generated. For rhizoid (Figure S5B), signal intensities were normalised so that the mean signal intensities of Venus and mScarletI are equal. For gemma epidermal cells, (Figure 2E), signal intensities were normalised so that the maximum signal intensities of Venus and mScarletI are equal.

### Morphometric analysis in Fiji

Height, width, and depth of fixed gemmae imaged on the confocal were measured in Fiji. To measure height and width, first, a maximum z-projection image of a gemma was rotated along the XY plane to orient the gemma stalk attachment point to the base of the image. Then the area selection tool was used to draw the smallest rectangle encompassing the whole gemma. The height and width of this rectangle was taken as the height and width of the gemma. To measure gemma depth, the orthogonal view function was used to manually find the XZ plane with the thickest gemma cross section. Within this XZ plane image, the line tool was used to draw a vertical line along the thickest part of the gemma, connecting the two flattened epidermal surfaces of the gemma. The length of this line was taken as the gemma depth. The number of cell layers making up this thickest part of the gemma was taken as the maximum cell layer number for the gemma.

### Quantification of gemma epidermal surface curvature

The MorphoGraphX manual^29^ and methods by Kirchhelle et al., 2016 were used as guides to carry out the below analysis. TIF stacks were loaded into MorphoGraphX for 2.5D segmentation of gemma epidermal cells. To extract the surface topology of the whole tissue, the image was first filtered using Gaussian blur (radius 0.3μm), then the Edge Detect function (threshold: 7000) was performed. Holes in the resulting mask were filled. A mesh was created using the Marching Cubes Surface algorism with a cube size of 2μm. After smoothing and subdividing the mesh, the SR2200 cell wall signal was projected on to the mesh. To segment the gemma epidermal cells, cells were manually seeded based on the projected cell wall signal, then Watershed segmentation was performed. After correcting segmentation errors and refining the segmentation, average surface curvature for a neighbourhood with a radius of 20μm was computed for each cell. The radius of 20μm was determined empirically to give an appropriate approximation of the average cell surface curvature for most gemma epidermal cells but not for very small cells. Hence statistical analysis was restricted to cells with an area greater than 150μm^2^.

### Identification of *ROP* homologues

To identify *ROP* homologues in land plant species with published genome assemblies, the *M. polymorpha* ROP amino acid sequence was used as the query sequence in tBLASTn and BLASTp searches in online genome/CDS and protein databases, respectively. To identify *ROP* homologues in species with only published transcriptome assemblies, tBLASTn searches were performed in online transcriptome databases. When genome, CDS, protein, or transcriptome databases of interest were not available on online BLAST servers, the datasets were downloaded for a local BLAST search on BioEdit^52^. Instead of relying on an arbitrary BLAST E-value threshold, each hit was examined manually to assess homology. As well as manually inspecting sequences for the presence of conserved domains (annotated in Figure S3F), the presence of the RHO domain was evaluated using the PROSITE and SMART protein domain databases^53,54^.

### Statistical analysis

Data plotted on graphs and statistically analysed in R.

## ACKNOWLEDGMENTS

We thank the Haseloff Lab (University of Cambridge) for providing the OpenPlant kit and the plasma membrane reporter line^40^. We thank Eva-Sophie Wallner and Victoria Spencer for critical reading of the manuscript. We thank Katharina Jandrasits and Magdalena Mosiolek (GMI), Helen Prescott and Lida Chen (University of Oxford) for assistance in the lab. We thank Charlotte Kirchhelle and the GMI/IMBA/IMP BioOptics team for advice on confocal imaging and image analysis. We thank Molecular Biology Services, Media Lab, and Lab Support of GMI/IMBA/IMP and the VBCF Plant Sciences unit for their support. This work was funded by a Biotechnology and Biological Sciences Research Council Doctoral Training Partnership Scholarship (Grant No. BB/M011224/1) to H.M., and a European Research Council Advanced Grant DENOVO-P (Project No. 787613) to L.D.

## SUPPLEMENTAL INFORMATION

**Figure S1.**
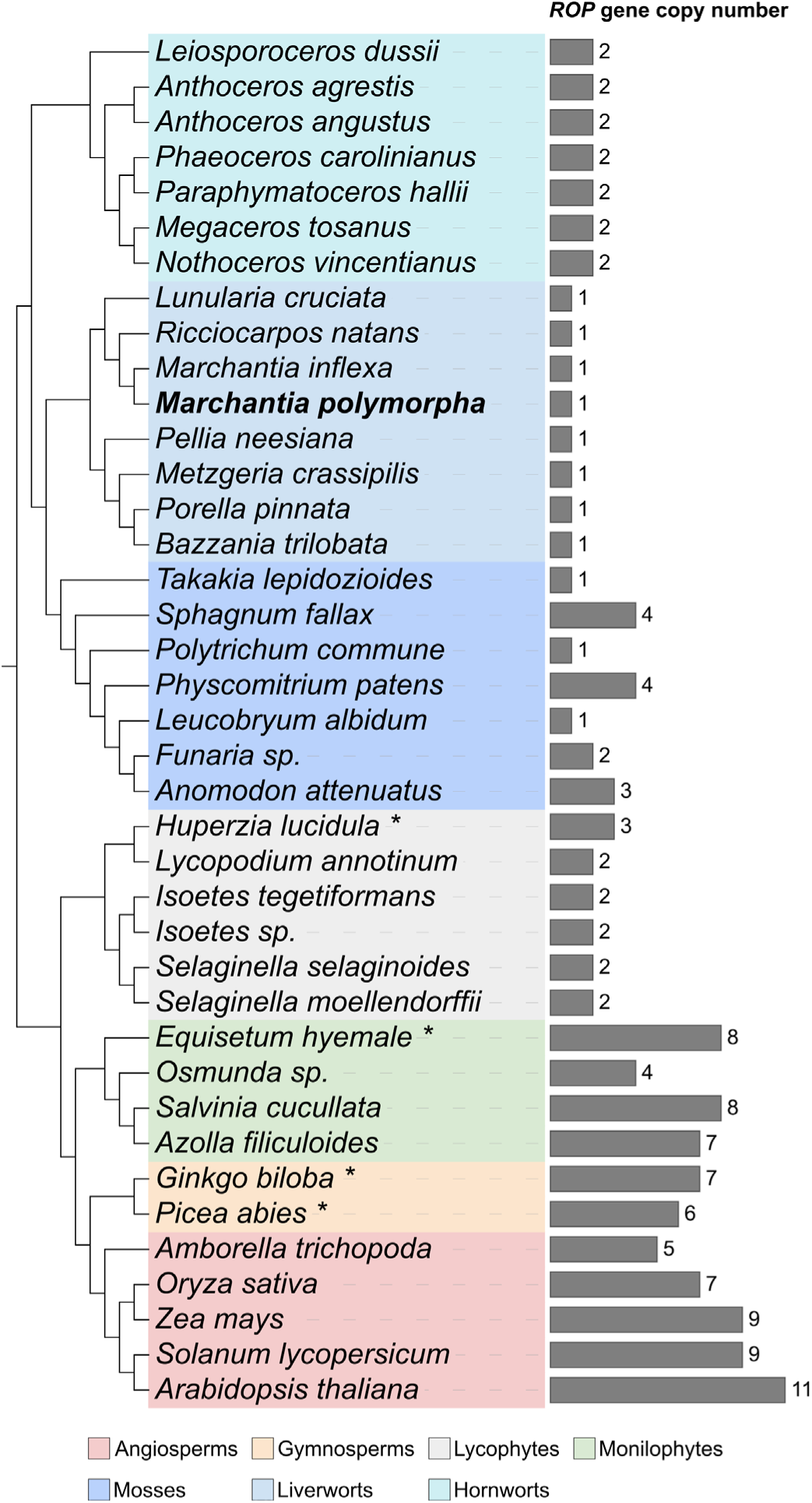
Liverworts, including Mar*chantia polymorpha*, encode only a single *ROP* gene in their genome, unlike most other land plants. Species tree of 39 land plant species combined with a bar graph displaying the number of *ROP* genes identified in the genome and/or transcriptome of each species. For four species (marked with an asterisk), the number of *ROP* genes encoded in their genome could not be determined with confidence due to poor quality genome/transcriptome assemblies. For these four species, the number of complete *ROP* genes identified with confidence are displayed and hence it is possible that these species encode more *ROP* genes. The species tree topology is based on ^55–57^.

**Figure S2.**
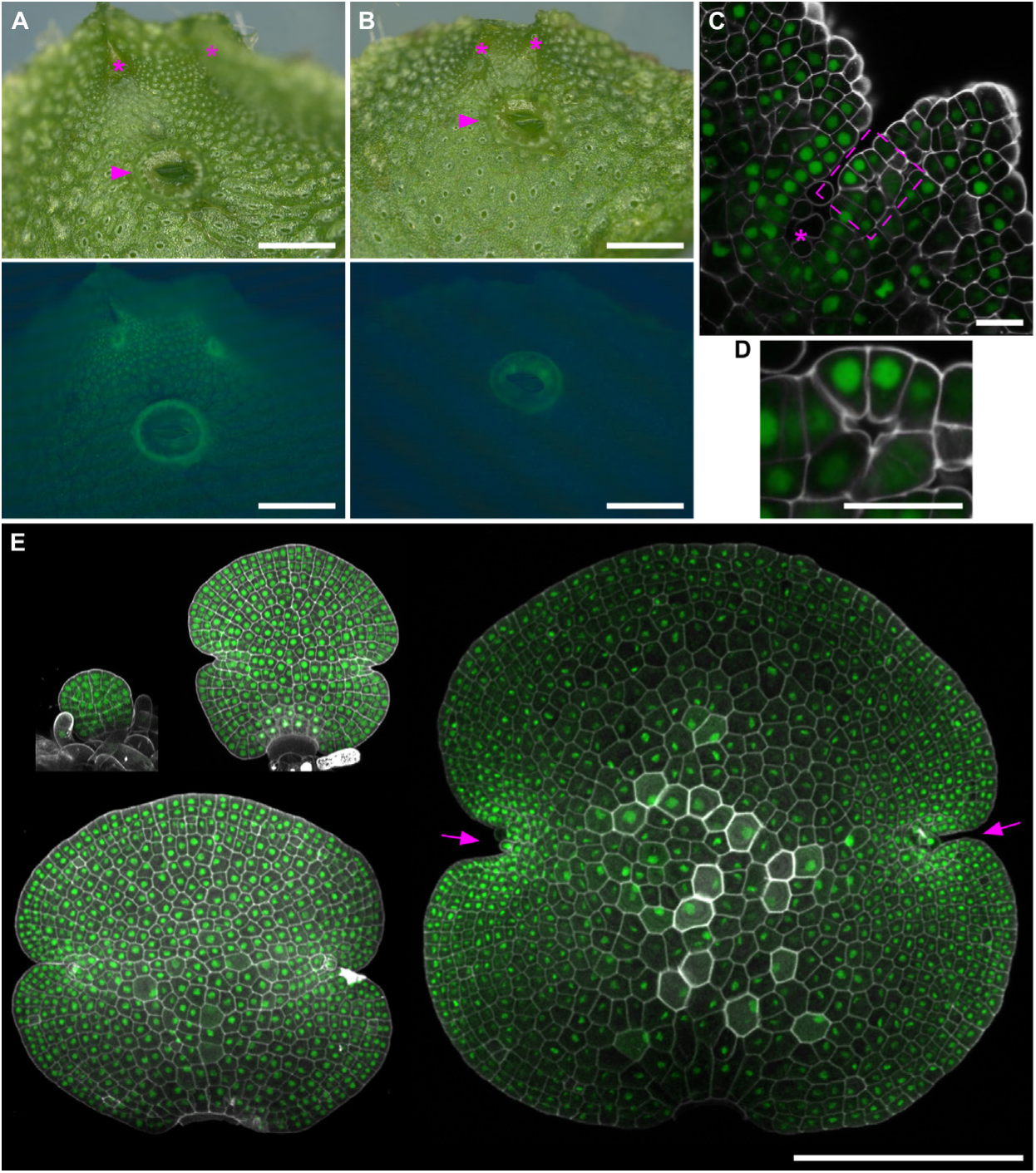
The Mp*ROP* gene is expressed ubiquitously throughout the thallus and gemma epidermis. Dorsal thallus epidermis of (A) *_pro_*Mp*ROP:NLS-Venus* and (B) wild-type (Tak1) imaged using a fluorescent stereomicroscope. Top row shows bright field images, with the meristematic notches marked with asterisks and gemma cups marked with arrowheads. Bottom row shows fluorescence images acquired through a YFP filter. Fluorescence from wild-type gemma cup is autofluorescence and hence fluorescence from gemma cup of the Mp*ROP* transcriptional reporter is also partially autofluorescence. Scale bar, 1mm. (C) The meristematic notch region of the *_pro_*Mp*ROP:NLS-Venus* line fixed, cleared, and stained with a cell wall dye (SR2200) before confocal imaging. Venus fluorescence is in green, and SR2200 fluorescence in grey. Asterisk marks the meristematic notch, and the box marks an immature air chamber. Cross section image was prepared in MorphoGraphX. Scale bar, 20μm (D) Higher magnification image of the immature air chamber marked in C. Scale bar, 20μm (E) Gemmae at different developmental stages (from top left to bottom right in acceding order of maturity) expressing *_pro_*Mp*ROP:NLS-Venus*. Gemma cell wall was stained with propidium iodide (PI) before confocal imaging. Venus fluorescence in green, PI fluorescence in grey. Arrows mark the meristematic notches. Scale bar, 200μm.

**Figure S3.**
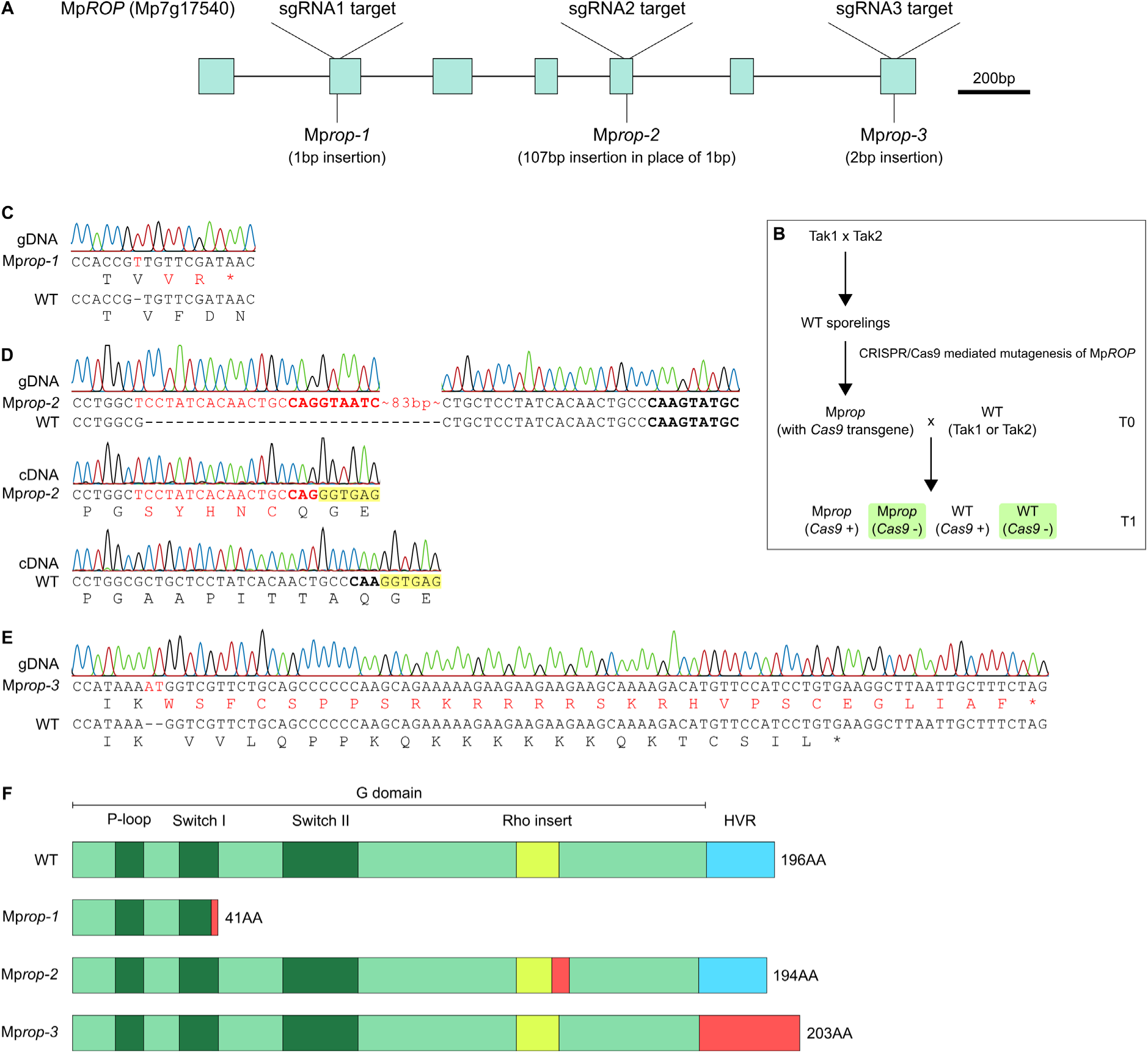
Mp*rop-1* and Mp*rop-3* are predicted to be total loss of function mutants and Mp*rop-2* is predicted to be a partial loss of function mutant. (A) Gene structure (from the start to the stop codon) of Mp*ROP*, annotated with target sites of sgRNAs used for CRISPR/Cas9 mediated mutagenesis. Mutation sites in Mp*rop-1*, Mp*rop-2*, and Mp*rop-3* are also marked. Turquoise boxes represent exons and lines represent introns. (B) *Cas9*-free Mp*rop-1*, Mp*rop-2*, Mp*rop-3*, and wild-type siblings (as controls) were isolated by crossing T0 Mp*rop* mutants to wild-type. (C-E) Genotyping through Sanger sequencing revealed specific mutations in the Mp*ROP* gene in Mp*rop-1,* Mp*rop-2*, and Mp*rop-3* alleles. Inserted nucleotides in the Mp*rop* alleles are in red. Predicted changes in amino acid sequence caused by the mutation are also in red. (D) A large insertion was found in the Mp*ROP* locus from Mp*rop-2* genomic DNA. The insert contains a sequence which closely matches the splice donor consensus sequence. This and the native splice donor sequence for intron 5 are in bold. Sanger sequencing of Mp*ROP* cDNA from Mp*rop-2* confirmed the sequence within the insert as the new splice donor site for intron 5. Nucleotides highlighted in yellow belong to exon 6. (F) MpROP protein domain architecture and predicted consequences of mutations in Mp*rop-1*, Mp*rop-2*, and Mp*rop-3* on the MpROP protein. Domains and motifs of interest are annotated. Domains with highly conserved sequences among RHO proteins are in dark green. HVR stands for Hypervariable Region. Amino acid sequence predicted to be altered in Mp*rop* mutants are in red. Annotations for P-loop and Rho insert is based on ^58^, and Switch I and Switch II are based on ^59^. G domain was determined using https://prosite.expasy.org/ and the region immediately following this was annotated as the HVR.

**Figure S4.**
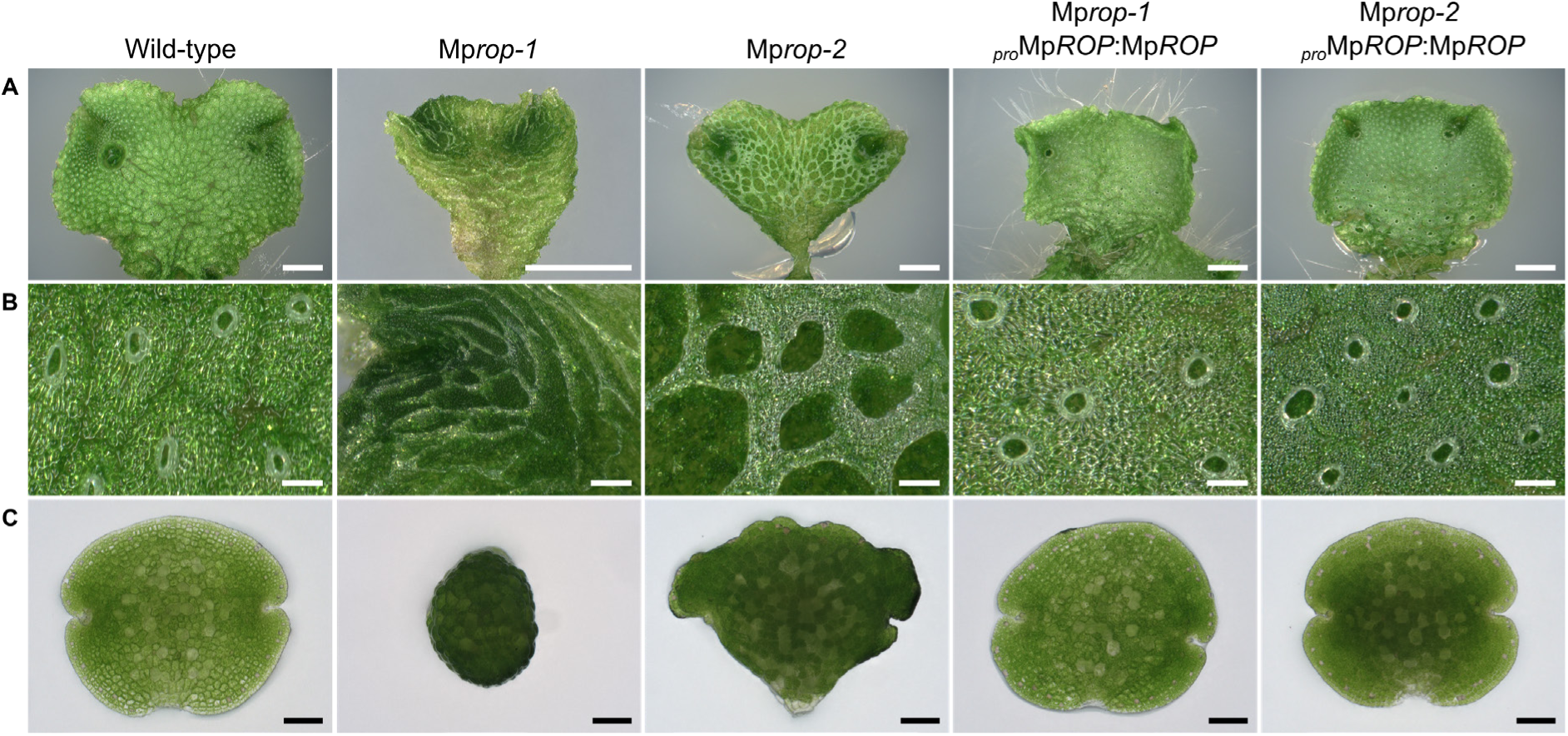
Genetic complementation confirms Mp*rop-1* and Mp*rop-2* are loss of function mutants. (A) Dorsal surface of a single thallus lobe of 14-day old gemmalings (scale bar, 2mm). (B) Dorsal thallus epidermis (scale bar, 200μm). (C) Gemma (scale bar, 100μm)

**Figure S5.**
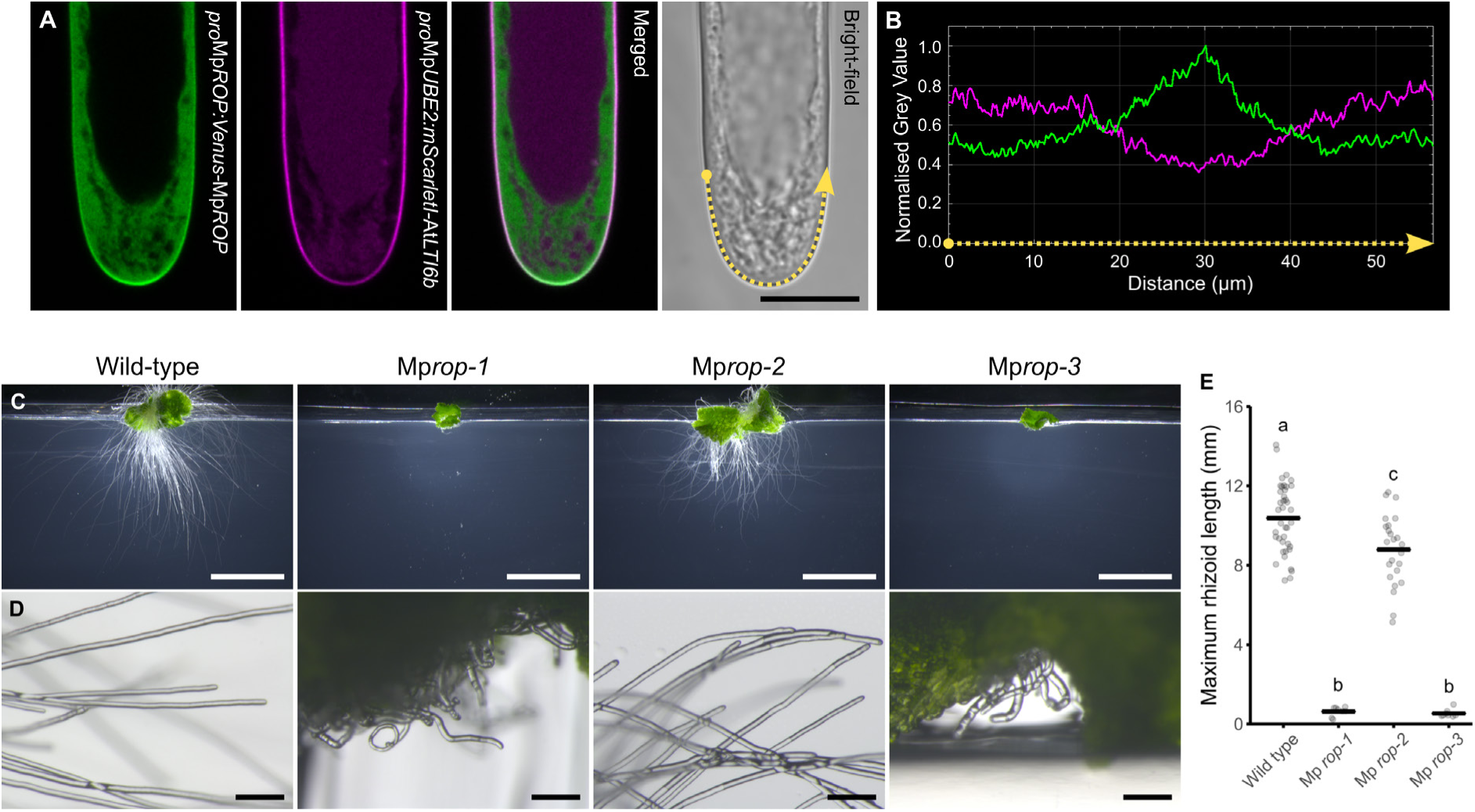
Venus-MpROP localises at the apex of tip growing rhizoid cells and Mp*ROP* is required for tip growth. (A) Confocal images of the growing end of a rhizoid cell expressing *_pro_*Mp*ROP:Venus-*Mp*ROP* and *_pro_*Mp*UBE2:mScarletI-*At*LTI6b* (plasma membrane reporter). Venus fluorescence is shown in green and mScarletI fluorescence in magenta. Dotted yellow line overlaid on the bright-field image marks the plasma membrane region along which fluorescence signal intensities from Venus-MpROP and mScarlet-AtLTI6b were compared. All images are Z-projections (sum slices) of five consecutive slices around the rhizoid medial plane. Scale bar, 20μm. (B) Normalised fluorescence intensities of Venus-MpROP (green) and mScarlet-AtLTI6b (magenta) along the plasma membrane region marked with the dotted yellow line overlaid in the bright field image in A. (C) Representative examples of wild-type and Mp*rop* mutants grown vertically on phytagel plates for 10 days from gemma. Scale bar, 5mm. (D) Representative rhizoid images of 7-day old gemmalings. Wild-type and Mp*rop-2* rhizoids are relatively straight. Mp*rop-1* and Mp*rop-3* rhizoids are very short and slightly curled. Scale bar, 200μm. (E) Rhizoids are significantly shorter in Mp*rop-1*, Mp*rop-2*, and Mp*rop-3* compared to in wild-type. Maximum rhizoid length was measured for 10-day-old gemmalings grown vertically on phytagel plates. Maximum rhizoid length was defined as the distance from the phytagel surface, on which the thallus sits, to the tip of the rhizoid which has grown the furthest away from the phytagel surface. On the graph, each grey circle represents the maximum rhizoid length of a different plant, and the black horizontal bar represents the mean maximum rhizoid length of all the plants sampled within one genotype. Different letters indicate groups with significantly different means, based on one-way ANOVA followed by Tukey’s HSD test (p < 0.05). The wild-type (n=40) sample consisted of four lines, two siblings of T_1_ Mp*rop-1*, and two siblings of T_1_ Mp*rop-2*. Samples for Mp*rop-1* (n=8), Mp*rop-2* (n=24), and Mp*rop-3* (n=8) each comprised two T_1_ lines.

**Table S1.**
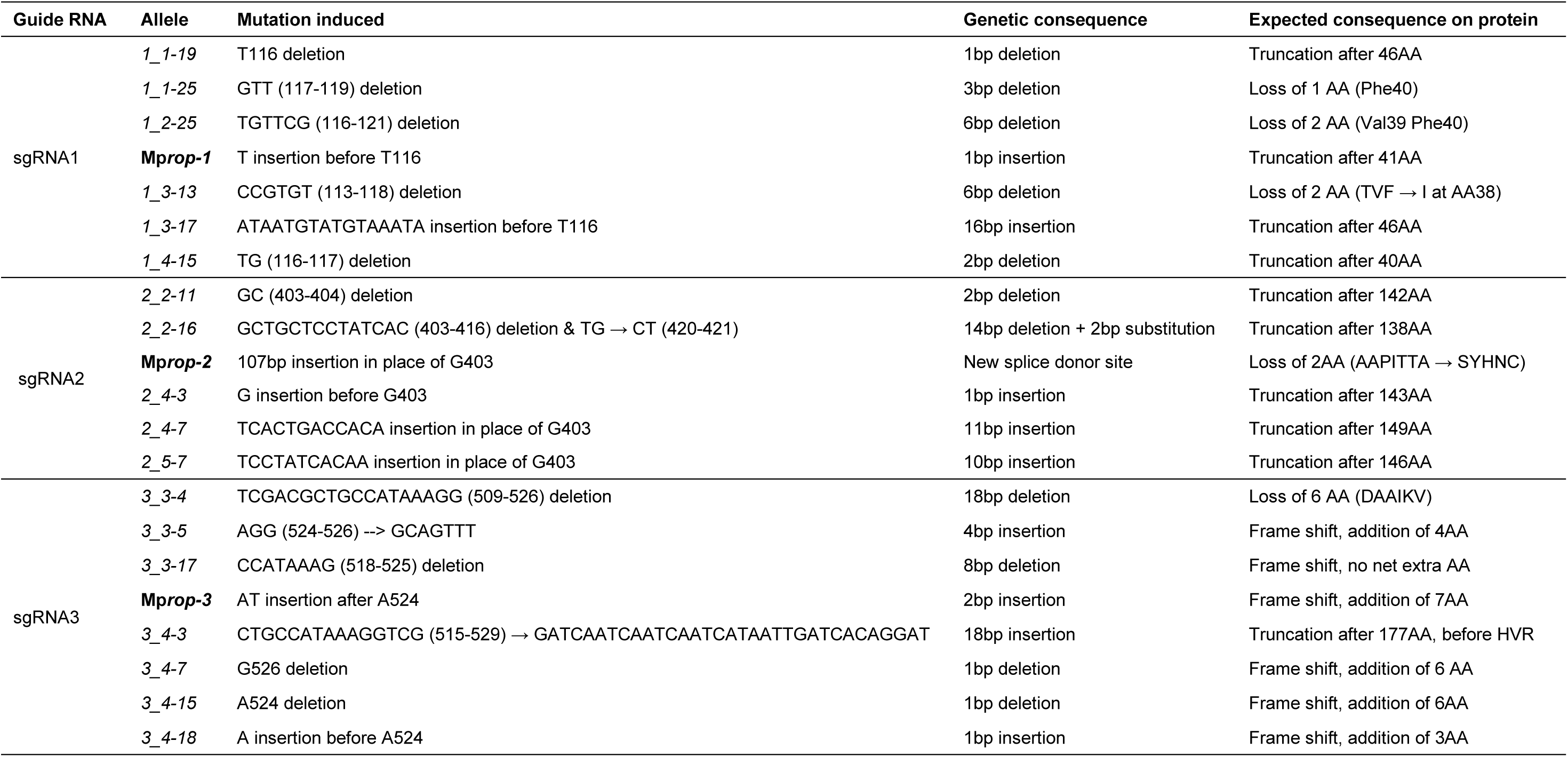
Twenty-one independent Mp*rop* mutants were generated using CRISPR/Cas-9. Three sgRNAs were designed to independently target three regions of the Mp*ROP* coding sequence. Sites targeted by each sgRNA are marked in Figure S3. Mutant alleles in bold are described in greater detail in Figure S3.

**Table S2.**
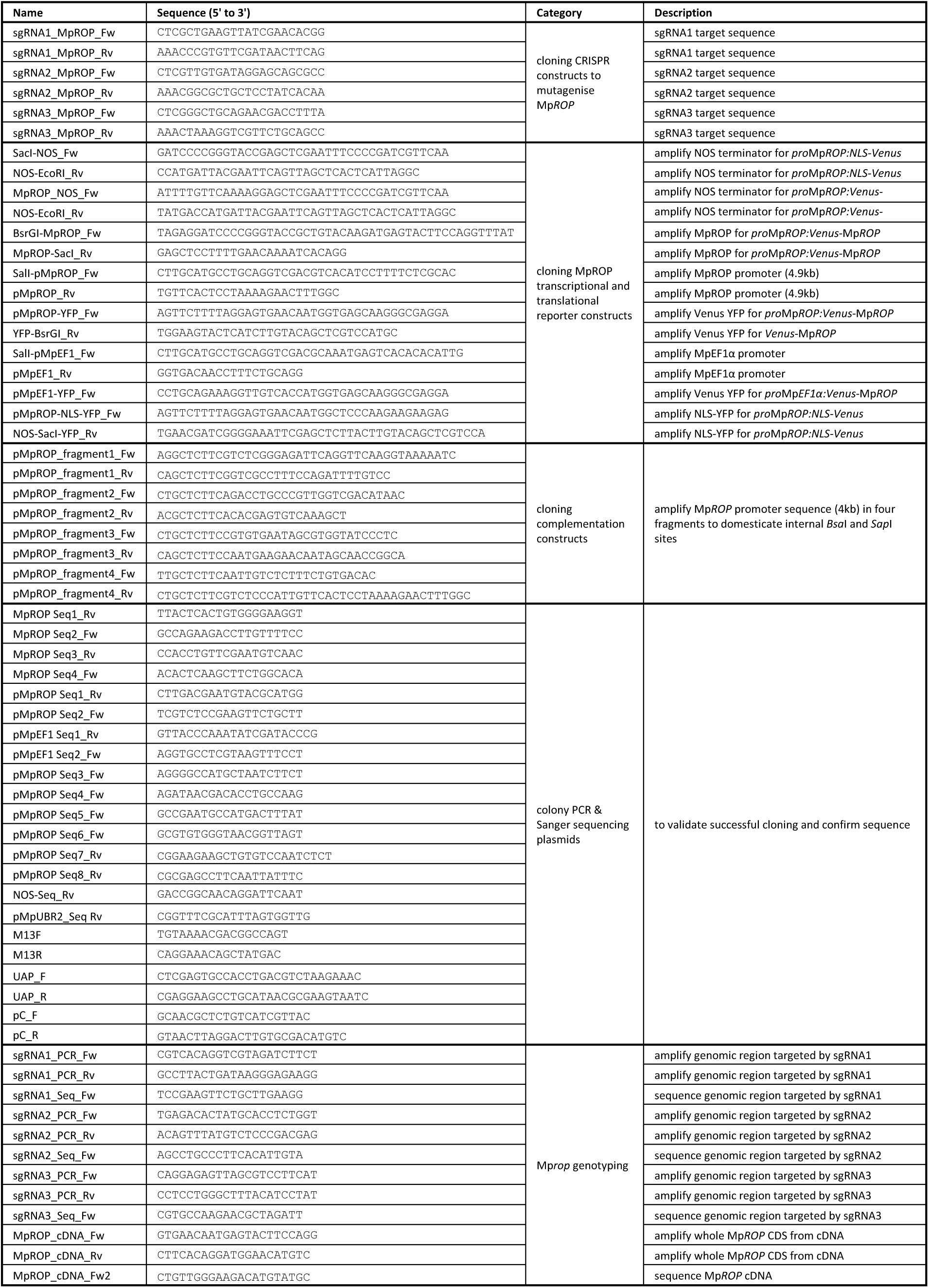
Primers used in this study.

